# A cell surface proteomic atlas reveals socioeconomic status associated immune diversity

**DOI:** 10.64898/2026.03.07.707513

**Authors:** Marouba Cisse, Wesley Huisman, Youvika Singh, Ibrahima Diallo, Moctar Gningue, Haiyu Wang, Alicia C. de Kroon, Rosanne A. M. Steenbergen, Geert H. Groeneveld, Babacar Mbengue, Maguette S. Niang, Tandakha N. Dieye, Alioune Dieye, Leendert A. Trouw, Souleymane Mboup, Kimberly Bonger, Maria Yazdanbakhsh, Moustapha Mbow, Simon P. Jochems

## Abstract

Socioeconomic status (SES) is a potent determinant of immune variation, yet unbiased approaches to holistically map the effect of SES on the immune system are lacking. We developed a high-dimensional flow cytometry-based profiling approach to analyse 331 cell surface proteins across 33 immune cell subsets within an SES-stratified Senegalese cohort, alongside a European cohort (Netherlands). We identified 108 SES-related markers across the immune system, revealing that lower SES individuals exhibited downregulation of surface proteins, affecting in particular adhesins, chemokine and complement receptors. Conversely, lower SES was associated with hallmarks of chronic activation and exhaustion including upregulation of immune checkpoints. Metabolic profiling demonstrated that while lower SES individuals displayed elevated baseline RNA transcription, higher SES individuals exhibited superior protein translation rates. We validated these SES-related immune trends in an independent cohort and provide an interactive online resource for exploring this surface protein atlas on immune cell subsets. Taken together, these findings provide a global overview of how the cell surface proteome varies by SES, and identify molecular changes that can affect vaccine efficacy and disease outcomes.

## INTRODUCTION

Inter-individual variation in the human immune system is a primary driver of global health disparities (Brodin & Davis, 2017). While genetics play a clear role in shaping the immune system, non-heritable influences, including environmental exposures and lifestyle factors account for nearly 58% of the divergence in immune response across populations (Brodin et al., 2015). This diversity in the immune system has effects on key clinical outcomes, including vaccine immunogenicity, immune responses to infections, allergic diseases, as well as cancer and responses to therapy (Abavisani et al., 2024; de Jong et al., 2021; Mbow et al., 2025; van Dorst et al., 2024). Within this environmental framework, Socioeconomic Status (SES) has emerged as a critical determinant of immune heterogeneity (Brodin & Davis, 2017; Snyder-Mackler et al., 2016), yet its molecular imprint remains largely unexplored.

SES has a significant influence on the immune system through a complex interplay of factors that includes urbanisation, lifestyle, environmental exposures, and diet (MacGillivray & Kollmann, 2014; Manurung et al., 2025; Mbow et al., 2014; Pyuza et al., 2024; Relman & Lipsitch, 2018; Rook, 2022). For instance, lower SES is associated with compromised control of persistent pathogens, such as cytomegalovirus (CMV), and elevated frequencies of pro-inflammatory immune cell subsets driven by epigenetic modifications (Azad et al., 2012; Dowd et al., 2009; Ravi et al., 2024; Stringhini et al., 2015; Tehranifar et al., 2013). Low SES also been shown to be associated with increased nasal colonization by pathobionts and reduced local immune responses in Indonesian schoolchildren (van Dorst et al., 2022).

Despite the epidemiological link between SES and immunity, the full extent of phenotypic changes related to SES remain unclear. Previous studies investigating SES-associated immune profiles have been constrained by limited parameters (Bertrand et al., 2024). As the functional interface between the immune cell and its environment, the surface proteome modulates cellular adhesion, migration, activation, and receptor-mediated signaling. Consequently, profiling these markers via flow cytometry provides a high-resolution readout for assessing the systemic impact of social determinants on the immune landscape (Ronsmans et al., 2022). Although high-throughput screening platforms have been established to discover immune cell surface using the LEGENDScreen™ kit (Liu et al., 2023; Magill et al., 2018), they are not typically used to compare between groups.

To address this gap, we performed high-dimensional profiling of over 300 cell surface proteins across 33 distinct immune cell subsets in cohorts from Europe (Netherlands) and West Africa (high and low SES, Senegal) using spectral flow cytometry. We identified, and validated a conserved SES-associated signature, characterized by the downregulation of critical surface proteins in lower SES individuals. We developed an interactive web application that can be used to explore this cell surface atlas. Within this setting of proteomic downregulation, we observed an uncoupling of transcriptional and translational efficiency along the SES strata. These findings establish a biological framework linking social determinants to fundamental cellular machinery, which may have implications in the disparities in vaccine effectiveness and disease outcomes.

## RESULTS

### Study population

To characterise immunological signatures associated with socioeconomic status (SES), we included healthy adult individuals from Europe and Senegal, matched for age and sex (Table S1). The Senegalese cohort was further stratified by SES based on living environments and socioeconomic criteria. The lower SES group displayed marked disparities in income security (100% irregular income, vs 38% in high SES, *P* = 0.026) and household density (average 3-fold higher occupancy, *P* < 0.001) compared to high SES (Table S2). While the groups were balanced for overall education level (*P* = 0.2), we observed a trend toward lower formal education in the low SES group education (0% reaching university, vs 50% in high SES). Moreover, limited healthcare accessibility (*P* = 0.018) and increased exposure to animals were characteristic of low SES (Table S2).

### Surface marker profiling uncovers SES-associated immune signatures

We developed a high-dimensional cell surface protein screening platform using spectral flow cytometry, based on the LEGENDScreen™ combined with sample barcoding and a backbone panel to identify cell subsets (Fig. 1A). Using a 23-parameter phenotyping panel, we identified 38 immune cell subsets via manual gating on which surface markers could be interrogated (Fig. S1 and Table S3).

**Fig 1.**
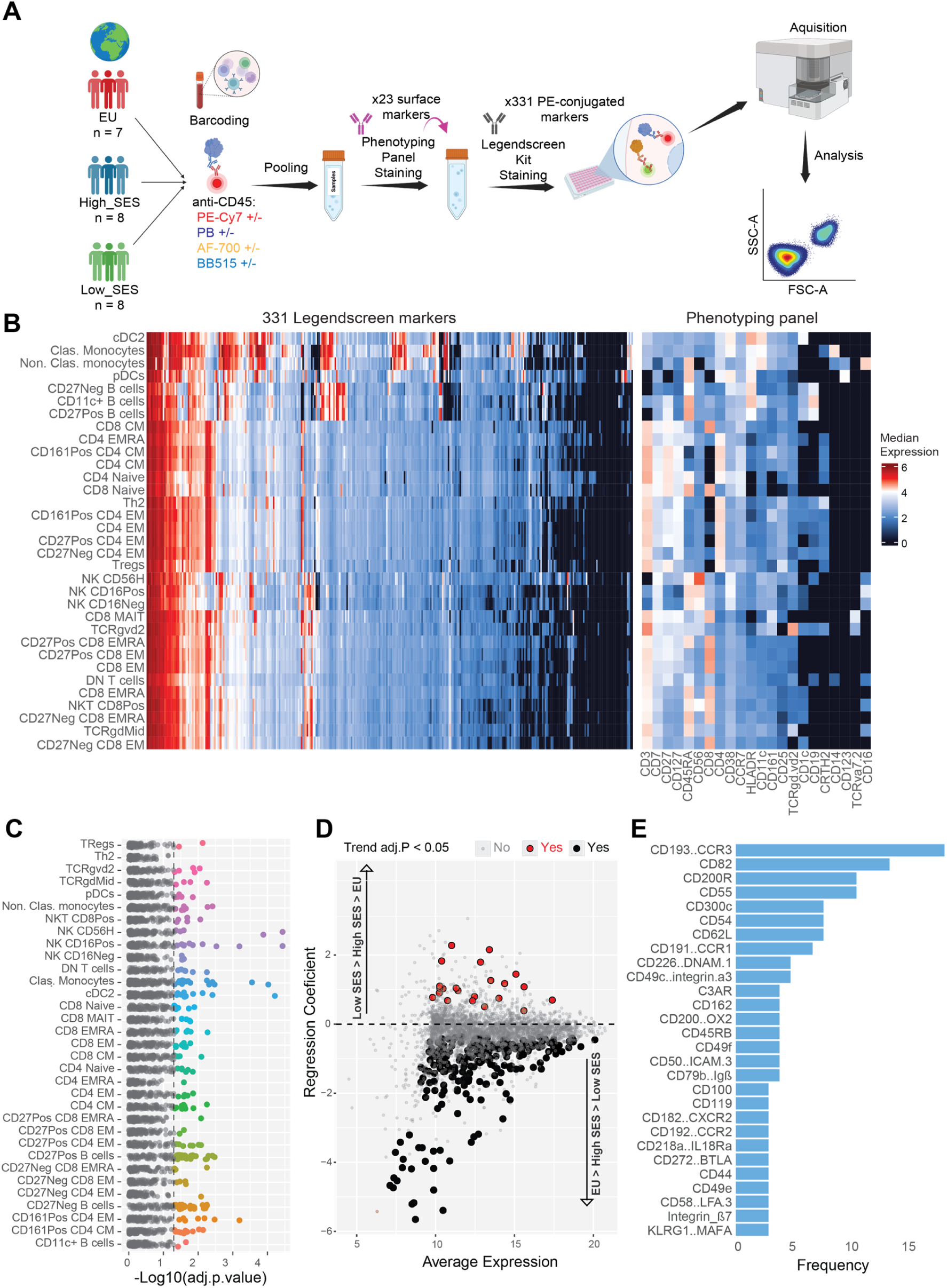
High-throughput immune profiling across socioeconomic populations. **A**) High-dimensional approach to map surface marker expression across immune cell subsets. **B**) Heatmap depicting the median expression of surface marker across manually gated immune cell subsets (y-axis). The left panel shows the screened surface proteins from the LEGENDScreen kit while the right ones are the subset defining markers from the phenotyping panel. **C**) Manhattan plot showing differentially expressed surface markers between groups across immune cell subsets (y-axis); dashed line indicates an FDR-adjusted P < 0.05 significance threshold. **D**) Linear mixed-effects model analysis comparing surface marker expression between SES groups following monotonic trend gradient (EU > Higher SES > Lower SES or the reverse). Black and red dots: FDR-adjusted *P* < 0.05; gray dots: non-significant. Y-axis shows regression coefficients indicating expression trend direction (positive: increased expression associated with lower SES; negative: increased expression associated with higher SES). **E**) Bar plots showing SES-related surface markers distribution across immune subsets.

First, we evaluated differential expression of cell surface markers across these subsets (Fig. 1B). To achieve this, we applied a limma-based trend analysis modelling groups as ordered factors to capture trends across multiple groups to test for a SES gradient from Europeans to higher SES Senegalese to lower SES Senegalese. Significant trends were followed by post-hoc pairwise comparison of all groups included in a linear mixed model, analogous to previous studies (Manurung et al., 2025).

We identified 108 unique surface markers across the 33 immune subsets that showed statistically significant SES-associated expression patterns (Fig. 1C). Of these, 91 surface markers were related to a higher SES across 32 immune subsets, whereas 17 were associated with lower SES (Fig. 1D) on 15 immune subsets. While differential expression was largely subset-specific, several markers displayed SES-associated changes across multiple cell types (Fig. 1E). Consequently, these 108 unique surface proteins corresponded to 263 significant marker-subset pairs, that is a marker that were SES-associated on a specific immune subset.

All 263 marker-subset pairs distinguished European from Senegalese individuals (Fig. 2A). Within the Senegalese cohort, 34 marker-subset pairs (13%) differed between high and low SES individuals, representing 21 unique surface markers across 16 immune subsets (Fig. 2B). For instance, the immune checkpoint CD160 (on CD8+ mucosal-associated invariant [MAIT] T cells), Integrin αM CD11b (on CD27 negative B cells) and the toll-like receptor-like molecule CD180 (TLR-4 homologue, classical monocytes) were upregulated in the low SES group. In contrast, marker such as Notch-2 (CD8+ central memory T [Tcm] cells), the cell adhesion molecule CD62L (classical monocytes and CD161+ CD4+ Tcm cells) and triggering receptor expressed on myeloid cells 1 (CD354-TREM1, cDC2) were associated with high SES.

**Fig 2.**
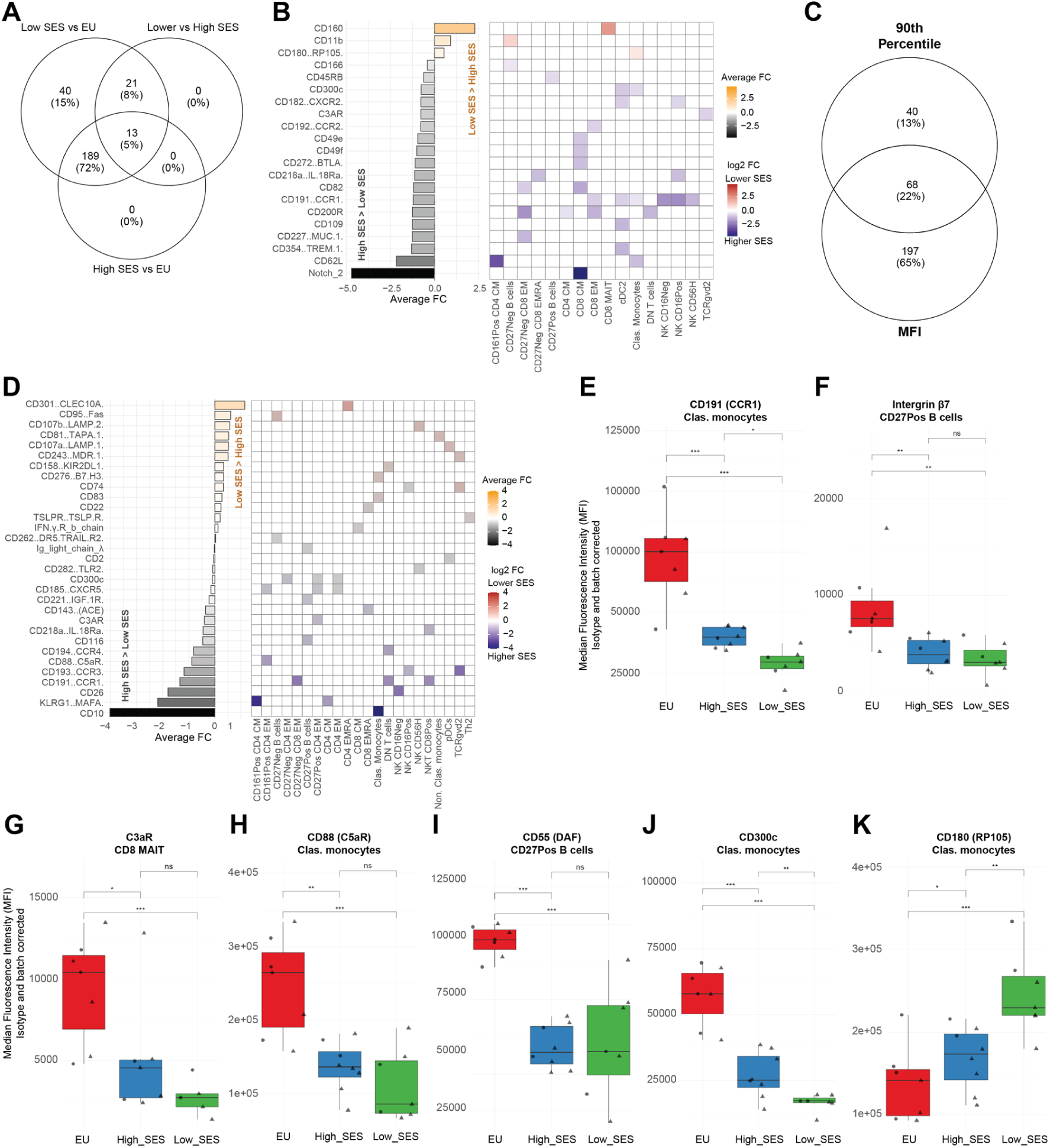
Socioeconomic status associates with differential expression of chemokine, complement, and costimulatory receptors. **A**) Venn diagram showing overlapping sets of cell surface markers that discriminated between socioeconomic groups. **B**) Immune markers differentiating Higher SES versus Lower SES individuals in Senegal along with immune cell subsets where they are differentially expressed using MFI data. **C)** Socioeconomic status-related immune markers that were differentially expressed in both MFI and 90^th^ percentile analyses (FDR-adjusted *P* < 0.05, trend analysis). **D)** Immune markers differentiating Higher SES versus Lower SES individuals in Senegal along with immune cell subsets where they are differentially expressed using 90^th^ percentile data. Examples of differentially expressed chemokine receptors between SES groups with CCR1 on classical monocytes (**E**) and Integrin B7 on CD27-B cells (**F**). Examples of differentially expressed complement receptors between SES groups with C3aR (**G**) on CD8+ MAIT, C5aR on classical monocytes (**H**) and the complement decay accelerating factor (DAF) on CD27+ B cells (**I**). Examples of differentially expressed costimulatory molecules and checkpoint receptors with CD200R on CD8+ EM (**J**), and CD180 on Clas. monocytes (**K**).

Given that activation markers often exhibit skewed distributions, with only a subset of cells expressing them, we performed parallel analyses using the 90th percentile (P90) of expression. This approach identified 108 SES-associated marker-subset pairs, corresponding to 58 unique surface markers across 29 immune subsets (Fig. 2C). Of these pairs 68 were consistent with the MFI analysis, while 40 pairs were exclusive to the P90 analysis (Fig. 2C), representing 31 unique surface markers across 22 subsets (Fig. 2D). For example, the angiotensin converting enzyme CD143-ACE (on CD8+ EMRA T [Temra] cells) was associated with higher SES while the Fas receptor CD95 (on CD27Neg B cells), and the immune checkpoint molecule CD276 (B7-H3) on classical monocytes were associated to lower SES (Fig. S3A-C).

To visualize immune marker expression across immune subsets and compare SES groups, we provided an interactive atlas, publicly accessible at https://cismar.shinyapps.io/LegendscreenShinyApp/. Each marker can be compared between groups, but also visualized between cell subsets to be able to identify marker expression patterns, with Figure 2E-K showing example markers extracted from the app. With this, we observed SES-dependent expression patterns for several surface markers essential for pathogen recognition, immune cell activation, and vaccine response.

For example, adhesion molecules and chemokine receptors, which regulate immune cell migration and positioning, were downregulated in lower SES individuals. This included reduced expression of chemokine receptor CCR1 (CD191) on classical monocytes (*P* < 0.001, Fig. 2E) and integrin β7 on CD27+ B cells (*P* < 0.05, Fig. 2F).

This downregulation pattern extended to complement anaphylatoxin receptors C3aR on CD8+ MAIT cells and C5aR (CD88) on classical monocytes, as well as the regulatory proteins decay accelerating factor (DAF/CD55) on CD27+ B cells (*P* < 0.05, Fig. 2G-I). To assess complement functionality, we measured serum C5b-9 deposition across classical, lectin, and alternative pathways (Fig. S4A-C). Complement activity was significantly increased in Senegalese individuals with higher SES in both alternative (*P* = 0.0232, Fig. S4D) and classical (*P* = 0.0243, Fig. S4E) pathways compared to lower SES individuals and Europeans. Conversely, Low SES Senegalese showed activity levels similar to Europeans with no significant differences in the lectin pathway (P = 0.2187, Fig. S4F). These findings indicate an uncoupling of complement activity and receptor expression on immune cells.

Costimulatory and checkpoint molecules that modulate T cell responses through balanced immune activation and regulation also showed SES-dependent variations. The activating receptor CD300c on classical monocytes (*P* < 0.001, Fig. 2J) decreased in low SES. However increased expression of CD180 (RP105), the structural homolog of Toll-like receptor 4 (TLR4) was related to lower SES (*P* < 0.05, Fig. 2K). Similarly, CTLA-4 on regulatory T (Tregs) cells showed a trend to increased expression in lower SES (*P* = 0.07 after multiple testing correction) (Fig. S3D).

### Immune subset variation related to SES

To validate key SES-associated markers, we included an independent cohort with 10 individuals per group (Table S5 and S6) that were analysed using three separate panels specific for T cells, myeloid cells, and B cells, respectively. We first assessed whether immune subsets varied by socioeconomic strata, by Integrating frequencies of manually gated immune cell subsets from individuals across LEGENDScreen discovery and the validation cohorts between groups. Sixteen immune cell subsets showed significant association with SES (Fig. 3A). Of these, five subsets were significantly enriched in a higher SES individuals (Fig. S5A), including CD8+ MAIT cells, basophils, and CD4+ CD8+ double-positive T cells (DP), Tregs, and classical monocytes. Conversely, eleven subsets were associated with lower SES (Fig. S5B) including CD11c+ B cells, Th2 cells, CD8+ NK T cells, and CD16-CD56dim NK cells with enrichment of differentiated effector memory phenotypes, specifically the CD27-CD8+ Temra, CD27-CD8+/CD4+ effector memory T (Tem) cell compartment.

**Fig 3.**
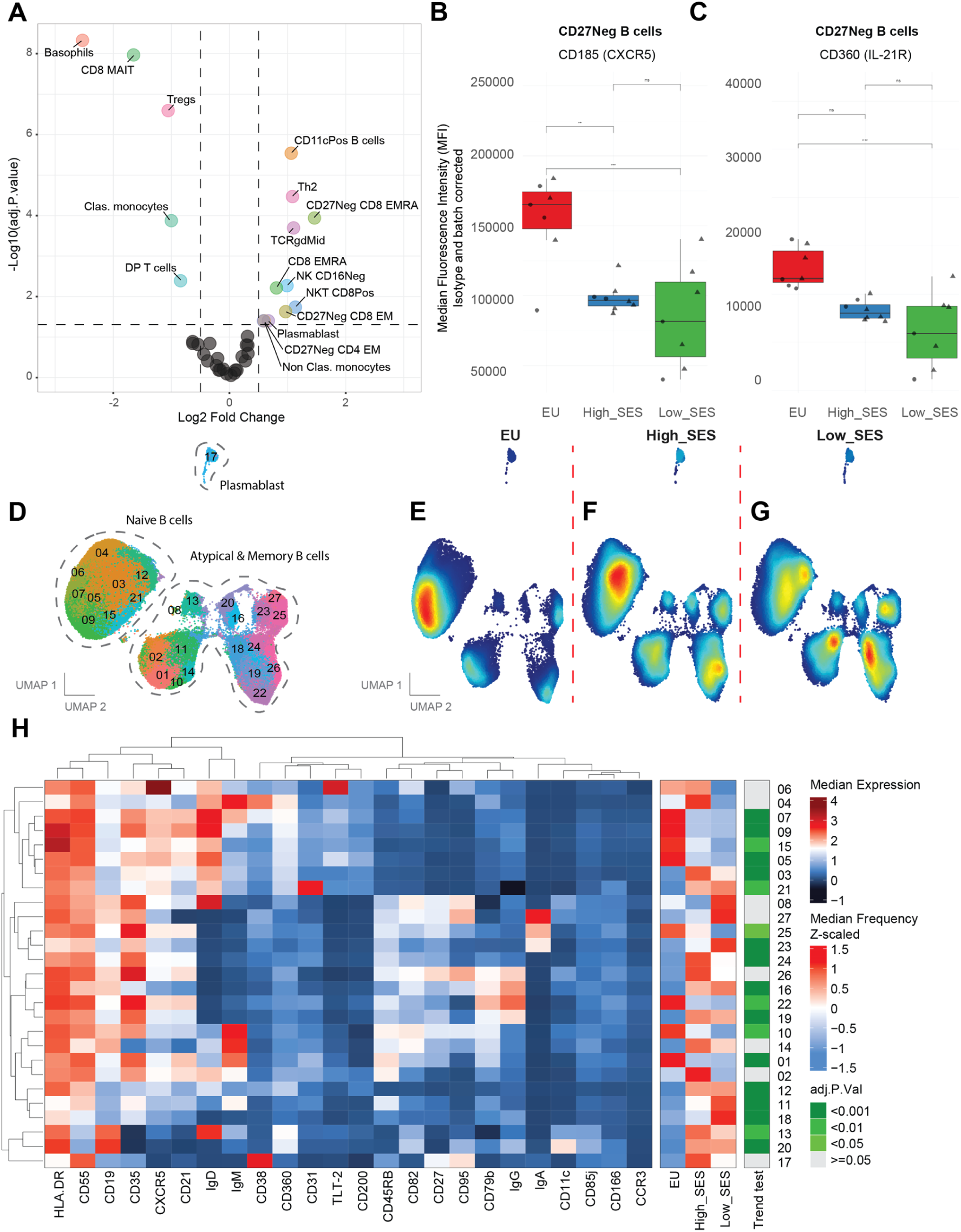
Germinal center-required markers decreased in low SES individuals. **A**) Volcano plot of frequency of immune cell subset differences between socioeconomic groups. Each point represents a cell subset. immune subsets that are differentially abundant (FDR-adjusted P < 0.05, trend analysis) with an at least Log2 fold-change differential frequency are indicated. Points below the horizontal dashed line denote cell subset frequencies without significant differences. **B-C)** Boxplot showing the average MFI of the chemokine receptor CXCR5 (CD185) and IL-21 receptor (CD360) between SES Groups on CD27-B cells. **D)** UMAP plot showing FlowSOM defined B cell clusters (n = 27) overlaid in distinct colors and differential subset abundancy between EU (**E**), Higher SES (**F**) and Lower SES (**G**). **H)** Heatmap depicting the median expression of phenotypic as well as SES-related immune markers across B cell clusters along with cluster median frequencies comparison between the SES Groups with a heatmap of significancy (FDR-adjusted P < 0.05).

### B cell proteins for germinal center involvement decreased in low SES individuals

Germinal center (GC) formation is a critical determinant of vaccine-induced humoral immunity. The chemokine receptor CXCR5 (CD185) directs B cells and T follicular helper (Tfh) cells to lymphoid follicles. Within GCs, IL-21 signaling through its receptor, IL-21R (CD360) on B cells promotes proliferation, somatic hypermutation, class-switch recombination, and differentiation into high-affinity plasma cells and memory B cells. Expression of CXCR5 (*P* = 0.0168, Fig. 3B) and IL-21R (*P* = 0.0298, Fig. 3C) on CD27-(naïve) B cells followed clear SES-related trend, with lowest expressions observed in individuals with a lower SES.

Among 52 markers differentially expressed on the three gated B cell subsets (CD11c+ atypical B cells, CD27+ memory and CD27-naïve), 25 (48%) were SES-related on both CD27+ and CD27-B cells, 18 (35%) were specific to CD27+ B cells, and 9 (17%) to CD27-B cells. The BCR component CD79b and the chemokine receptor CD193 (CCR3) were related to SES across all B cell subsets (Fig. S6A).

To validate differentially abundant proteins in additional individuals, and assess co-expression of these markers, we developed a targeted B cell panel (Table S7), focusing on functionally relevant SES-associated markers. This confirmed SES-dependent expression patterns for 15 unique surface proteins across the B cell subsets (23 marker-subset pairs), including germinal center markers CXCR5 and CD360 within the validation cohorts (Fig. S7A-B).

Manually gated B cell populations are overlaid in the UMAP dimension shown in Supplementary Fig. 7B, and unsupervised clustering using FlowSOM identified 27 B cell clusters (Fig. 3D). B cells were differentially distribution among Europeans (Fig. 3E), Senegalese with a higher SES (Fig. 3F) and those with lower SES (Fig. 3G), showing a clear gradient from Europeans to high SES to low SES Senegalese. Comparison of cluster frequencies across the SES gradient revealed that 19 out of 27 B cell clusters differed significantly between groups, confirming the discriminatory capacity of selected markers based on the initial legendscreen (Fig. 3H). Several naïve B cell clusters expressing CD360 (IL-21R) and CD185 (CXCR5) were indeed significantly enriched in higher SES individuals, clusters 05 (also TLT-2+ CD200+), cluster 07 (CD38+ TLT-2+), and cluster 09 (TLT-2^low^)(Fig. S6C). Conversely, several IgG+ clusters were associated with lower SES, in particular with expression of tetraspanin CD82 and activation marker/death receptor CD95: cluster 16 (CD82+ CD95+), cluster 18: (CD21^low^ CD35+), cluster 19: (CD21+ CD82+), and cluster 20 (CD95+ CD11c+)(Fig. S6D).

These findings indicate that lower SES is associated with downregulation of germinal center required surface proteins in naïve B cell compartment, with increased clusters of IgG+ B cells expressing activation markers.

### Myeloid compartment surface markers reveal contrasting allergic and inflammatory programming

Next, we examined the expression of cell surface markers on myeloid cell subsets within the discovery cohort. FcεRIα, the high-affinity IgE receptor, showed increased expression on classical monocytes with higher SES (*P* = 0.0259) (Fig. 4A). Conversely, increased expression of CD107b (*P* = 0.0291), the lysosomal-associated membrane protein 2 (LAMP-2) was related to lower SES (Fig. 4B).

**Fig 4.**
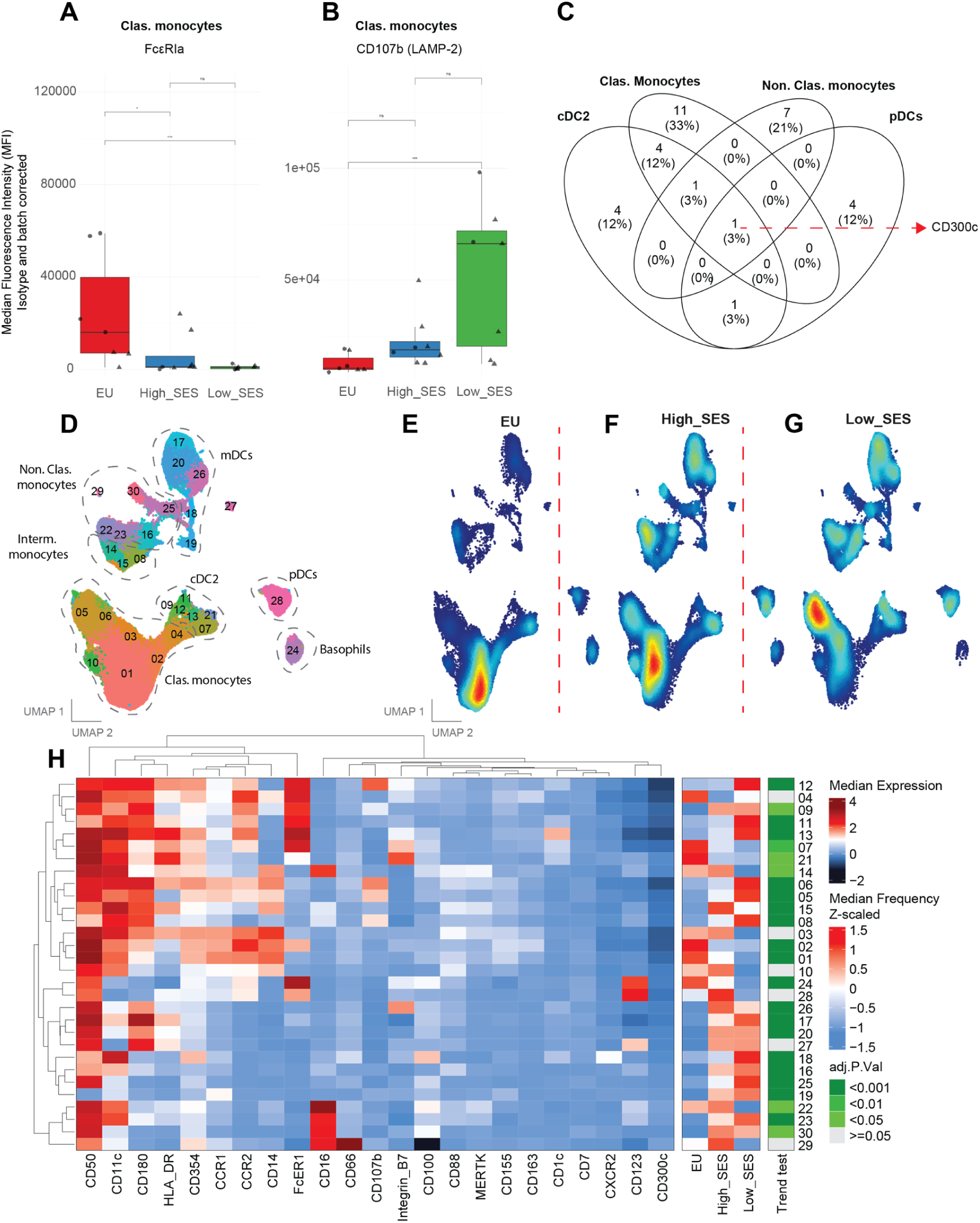
Surface markers reveal contrasting allergic and inflammatory programming in myeloid subsets. **A-B)** Boxplot showing the MFI expression of the high-affinity IgE Fc receptor (FcεRIα) and lysosomal-associated membrane protein 2 (LAMP-2) (CD107b) between SES Groups on classical monocytes. **C)** Venn diagram showing overlapping sets of surface markers differentially expressed across myeloid subsets. **D)** UMAP plot showing FlowSOM defined myeloid cell clusters (n = 30) overlaid in distinct colors. Density plot showing differentially abundant immune cell clusters between EU (**E**), Higher SES (**F**) and Lower SES (**G**). **H)** Heatmap depicting the median expression of phenotypic as well as SES-related immune markers across myeloid clusters along with cluster median frequencies comparison between the SES Groups with a heatmap of significancy (FDR-adjusted P < 0.05).

Comparing the 33 differentially expressed proteins across myeloid subsets, immune checkpoint protein CD200 was SES-related on pDCs and cDC2, and the chemokine receptor CCR1 was differentially expressed on all myeloid subsets except pDCs. Notably, CD300c, an activating receptor regulating inflammatory responses, was expressed in a SES-associated pattern across all myeloid subsets. Most markers were immune subset-specific: 11 markers (33%) were SES-associated in classical monocytes, 7 (21%) in non-classical monocytes, 4 (12%) in pDCs, and 4 (12%) in cDC2 (Fig. 4C).

We validated 13 SES-associated marker-subset pairs, representing 15 unique surface proteins across the myeloid subsets in the independent cohort (Table S8 and Fig. S8A). The inflammatory phenotype related to lower SES characterized by increased expression of LAMP-2 remain consistent across both the discovery cohort (Fig. 4B) and the validation cohort (Fig. S7C).

Unbiased FlowSOM clustering identified 30 distinct myeloid clusters based on this panel (Fig. 4D), which exhibited differential distribution among Europeans (UMAP, Fig. 4E), higher SES Senegalese (Fig. 4F), and lower SES Senegalese participants (Fig. 4G). Gradient analysis revealed significant frequency differences for 24 out of 30 myeloid clusters between SES groups, confirming that identified SES-related markers can discriminate between SES groups (Fig. 4H). Individuals with high SES were characterised by significant enrichment of myeloid clusters expressing FcεRIα, the immune checkpoint ligand CD155 (a TIGIT co-receptor) and the signalling protein CD100/SEMA4D, cluster 02 (CD163+ CD155+) and cluster 07 (CD100+ Integrin B7+)(Fig. S8B). In contrast, CD107b+ clusters were enriched in individuals with lower SES, specifically cluster 05 (HLA-DR^low^) and cluster 06 (MERTK+)(Fig. S8C).

In summary, our findings demonstrate that SES drives distinct myeloid reprogramming, with individuals with higher SES exhibiting an allergy-related profile while lower SES associated with pro-inflammatory phenotype.

### T cell immune checkpoint expression correlates with socioeconomic status

Focusing on T cells demonstrated that immune checkpoint expression on T cells associated significantly with SES. Individuals from higher SES backgrounds exhibited increased CD200R expression on CD8+ Tem cells (*P* = 0.0231, Fig. 5A) and elevated CD272 (BTLA) expression on CD8+ Tcm cells (*P* = 0. 0368, Fig. 5B). In contrast, lower SES associated with increased CD57 expression on Tregs (*P* = 0. 0356, Fig. 5C) and elevated CD160 expression on CD8+ MAIT cells (*P* = 0. 0298, Fig. 6D). CD200R (forms axis with CD200-OX2) as well as BTLA (forms axis with HVEM-CD270) function as co-inhibitory receptors that suppress immune activation (Derre et al., 2010), while CD160 regulates CD8+ T cell function and has emerged as a target to overcome anti-PD-1 resistance in cancer immunotherapy (Zheng et al., 2025). CD57 marks terminally differentiated T cells with enhanced cytotoxic potential (Brenchley et al., 2003).

**Fig 5.**
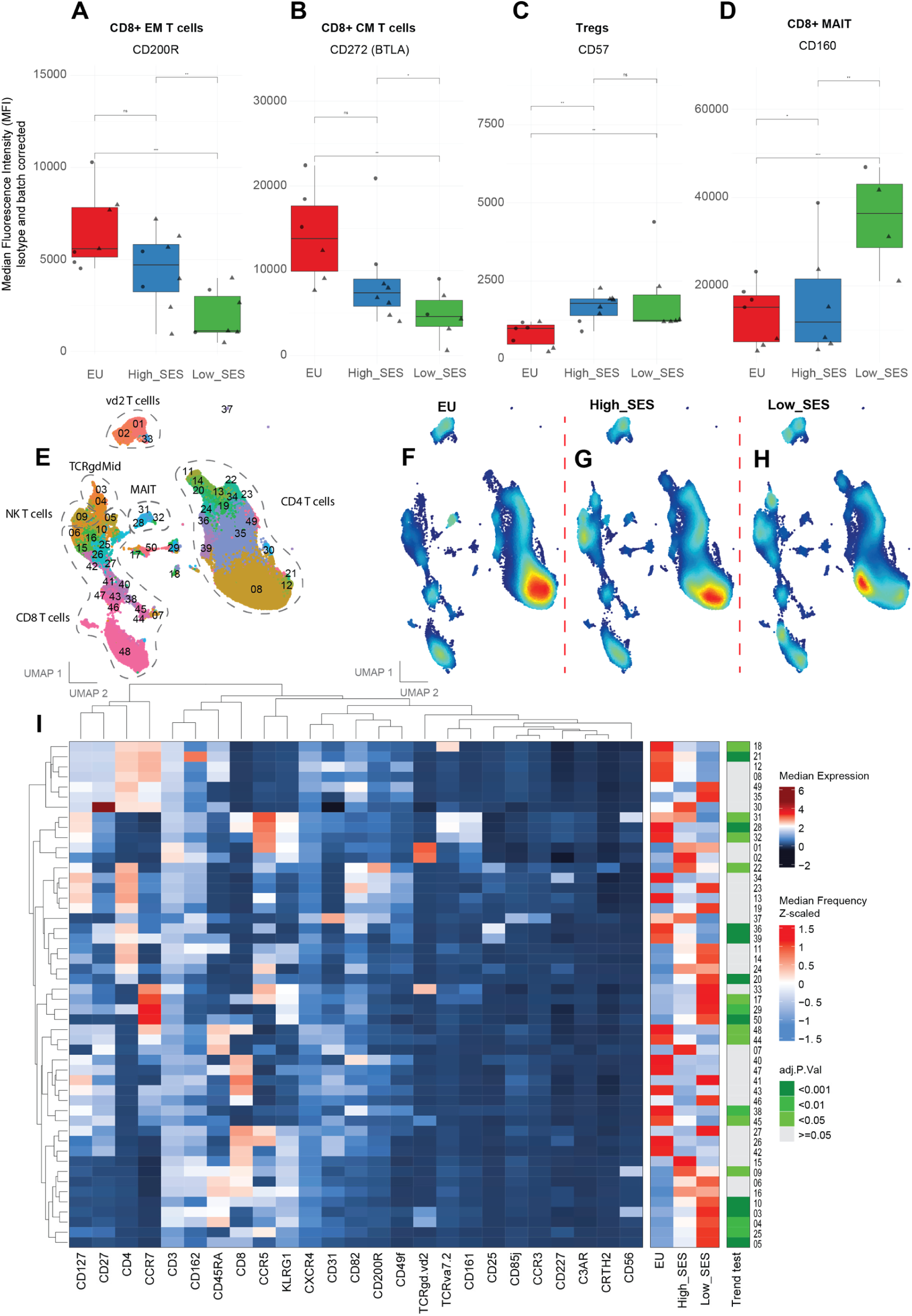
Immune checkpoint expression correlates with socioeconomic status. **A-D)** Boxplot showing the MFI expression of the immune checkpoint CD200R (on CD8+ EM T cells), CD272 (on CD8+ CM T cells) as well as CD57 (on regulatory T cells) and CD160 (on CD8+ MAIT cells). **E)** UMAP plot showing FlowSOM defined T cell clusters (n = 50) in distinct colors and differential subset abundancy between SES Groups (**F**-**H**). **I)** Heatmap depicting the median expression marker of T cell clusters and z-score cluster frequencies comparison between the SES Groups with a FDR-corrected p-value less than 0.05 considered significant.

**Fig 6.**
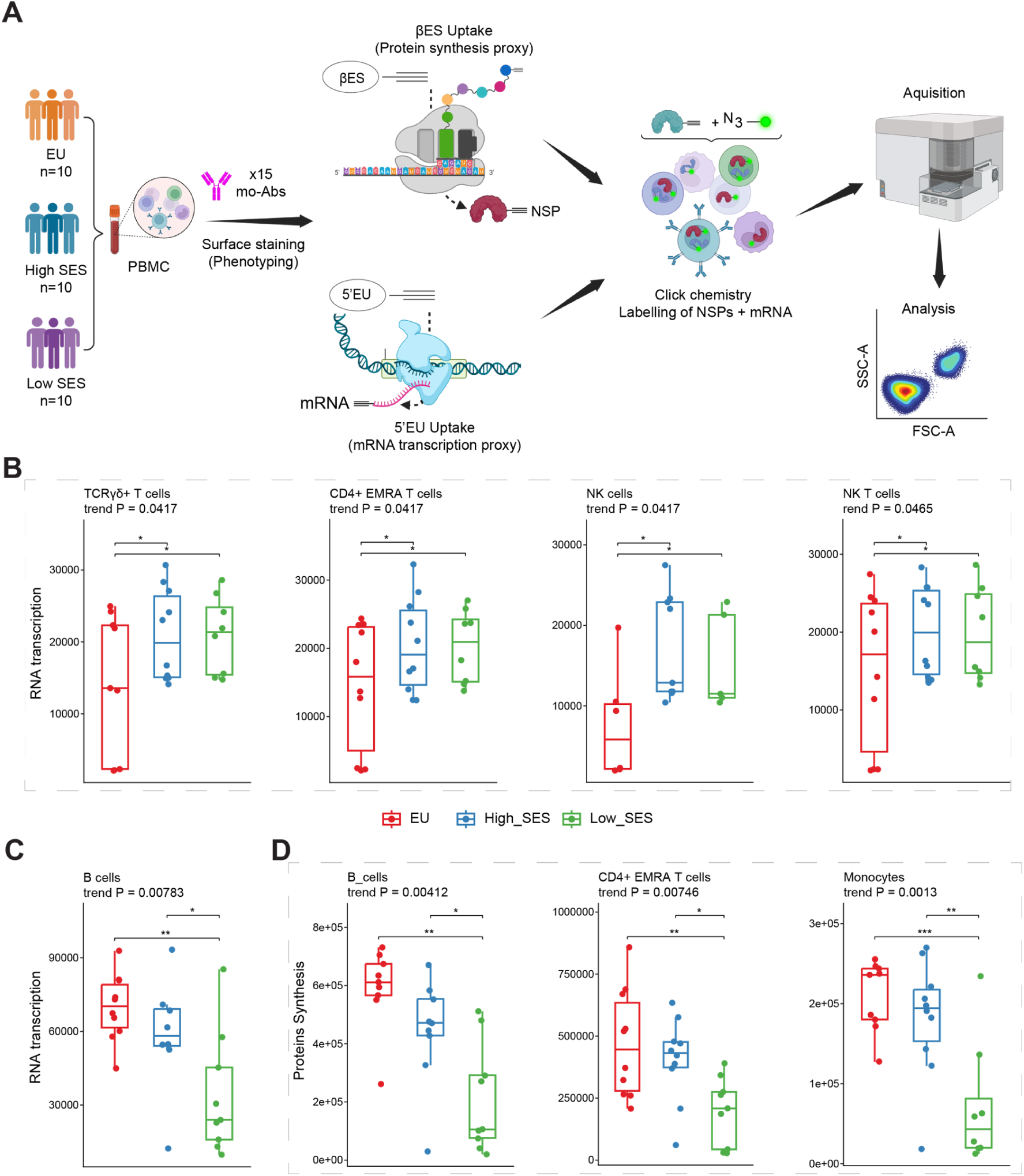
Cellular metabolism rates demonstrate SES-dependent regulation in activated immune cells. **A**) Spectral flow cytometry quantification of 5’EU and βES incorporation in PBMC after 1 hour incubation with 500 µM of βES or 5’EU in RPMI 10% hi-FCS to assess changes in transcriptional and translational rates across immune subsets between groups. Where indicated, stimulation was done overnight (16 hours) using a mixture of CPG-ODN2006 (1 µM), LPS (100 ng/mL), aCD3 (1 µg/mL), and aCD28 (10 µg/mL) in RPMI 10% hi-FCS. Boxplots of fluorescence intensity of bioorthogonal analogues incorporation across socioeconomic status (SES) groups, relative to 5’EU uptake (RNA transcription) in resting (**B**) and activated (**C**) immune cell subsets or protein synthesis (βES uptake) in activated immune cells (**D**).

Of these SES-associated markers, 10 unique surface proteins across the T cell subsets (17 marker-subset pairs) were validated in a the independent cohort using a targeted T cell panel (Table S9 and Fig. S9A). Consistent with our initial findings, we observed enrichment of T cell clusters expressing immune checkpoint proteins showing SES-dependant changes between the groups. For example, CD200R expression on CD8+ Tem cells on showed significant SES-associated changes mirroring the discovery cohort (Fig. S7D).

Unsupervised FlowSOM clustering identified 50 distinct T cell clusters (Fig. 5E), which were differentially abundant among European (Fig. 5F), higher SES Senegalese (Fig. 5G), and lower SES Senegalese (Fig. 5H). Trend analysis revealed significant frequency differences for 23 out of 50 clusters between SES groups (Fig. 5I). T cell clusters expressing CD200R were indeed enriched in higher SES individuals, specifically cluster 38 (CD49f+ CD82+)(Fig. S9B). Conversely, KLRG1+ clusters were enriched in individuals with lower SES, particularly those expressing the adhesion/chemokine receptors CCR5 such as cluster 05 (CCR5-), cluster 10 (CCR5+), and cluster 25 (CD31+ CCR5+ CXCR4+)(Fig. S9C).

Taken together, lower SES associated with significant alterations in the T cell compartment characterised by enrichment of terminally differentiated subsets expressing exhaustion and senescence markers.

### Cellular metabolism rates demonstrate SES-dependent regulation in activated immune cells

Given the general trend toward lower surface protein expression with lower SES (Fig 1D), we investigated whether protein translation and mRNA transcription rates varied across the SES gradient. We measured 5-ethynyl uridine (5’EU) and β-ethynylserine (βES) incorporation, as a readouts for RNA transcription and protein synthesis, respectively, across immune cell subsets in PBMCs from age- and sex-matched participants (n=10/group, Fig. 6A). Validation experiments confirmed that 5’EU and βES specifically labelled nascent mRNA and proteins, respectively, as incorporation was completely abrogated by the transcription inhibitor Actinomycin D and the translation inhibitor Cycloheximide respectively (Fig. S10A-B).

Metabolic profiling revealed an uncoupling of transcription and translation associated with SES. At baseline, lower SES was associated with increased RNA transcription in TCRγδ+ T cells, CD4+ Temra cells, NK cells and NK T cells (Fig. 6B). However, this elevated transcriptional baseline did not result in increased protein output, as ex vivo protein synthesis rates did not differ between SES groups.

Furthermore, lower SES exhibited reduced mRNA induction following overnight TLR, TCR, and BCR co-stimulation with differential transcription rates observed in B cells (Fig. 6C). Consistent with this blunted transcriptional response, protein synthesis rates following stimulation were highest in Individuals with high SES and lowest in those with low SES, particularly in stimulated B cells, CD4+ Temra cells and monocytes (Fig. 6D).

Together, these findings suggest that lower SES is characterized by reduced protein translational efficiency despite elevated mRNA transcriptional rates at baseline which seems to associate with lower responsiveness upon activation.

## DISCUSSION

In this study, we established a high-throughput approach to characterize immune signatures associated with socioeconomic status (SES). By profiling over 300 surface markers across 33 immune subsets, we demonstrate that SES is not only a demographic variable but a potent biological determinant that fundamentally reshapes immune system architecture in healthy adults. To facilitate exploration of this dataset by others, we have deployed an interactive webtool that can be freely accessed to interrogate markers of interest. While previous studies have noted differential immune profiles between rural and urban populations (Manurung et al., 2025; Mbow et al., 2014; Pyuza et al., 2024). These studies were often limited to discrete geographic comparisons or pre-defined parameters. Furthermore, the lack of large atlas-type datasets from global South is expected to drive future inequalities in health care and vaccine responses (Amoah et al., 2025). Here, we reveal that socioeconomic gradients systemically remodel the immune landscape, characterized by a specific downregulation of surface receptors and a significant metabolic reprogramming.

Individuals from lower SES backgrounds displayed a signature of chronic immune activation and exhaustion. We observed an expansion of effector memory T cells and NK subsets, coupled with upregulation of senescence-associated markers (CD57, CD160) and inhibitory checkpoints (CTLA-4 on regulatory T cells). Similarly, IgG+ B cells, including CD11c+ atypical B cells, expressing the receptor CD95 (FAS) were increased in lower SES individuals. These phenotypes likely reflects cumulative history of pathogen burden driven by higher household density, zoonotic contact and environmental exposures to micro-organisms, resulting in the enrichment of terminal differentiation of the adaptive compartment (Brenchley et al., 2003; Klopack et al., 2022). While this immune status may offer heightened protection against immediate threats but accelerated immune system aging, potentially impairing immune response to persistent pathogens like CMV and diminish effective cancer immunosurveillance (Dowd et al., 2009; Pawelec, 2017).

Crucially, we observed a significant downregulation of CXCR5 and IL-21R on B cells from lower SES individuals. These markers are essential for germinal center formation in vaccine-induced immunity. This alteration suggests a structural impairment in the ability to generate high-affinity antibodies and durable immunity (Kim et al., 2022; Kuchen et al., 2007; Linterman et al., 2010; Zotos et al., 2010). This aligns with observations that vaccine immunogenicity, mostly measured by antibody responses is often lower in low-income populations (Abavisani et al., 2024). The hypothesis that vaccine hyporesponsiveness is driven by altered migration patterns in lymphoid tissues and reduced germinal center engagement could be tested by employing fine needle aspiration of draining lymph nodes at the site of vaccination in global south populations (Turner et al., 2020). Such insights could allow vaccination strategies to be optimized for socioeconomic contexts through improved adjuvant selection and modified dosing schedules (Kurtovic & Beeson, 2021).

We also, observed population-specific dynamics in complement receptors and regulators. Despite the downregulation of surface complement receptors (C3aR and C5aR) across various immune cell subsets in low SES Senegalese individuals, complement activity was paradoxically elevated in Senegalese with higher SES compared to low SES Senegalese and Europeans. This uncoupling may reflect in vivo activation by subclinical exposures absent in European settings or driven by genetic polymorphisms, which were not assessed in this study design. Alternatively, diminished receptor expression could trigger compensatory upregulation of complement function. Integrating longitudinal sampling with infectious disease surveillance is necessary to clarify these relationships, as previously described (Dunkelberger & Song, 2010; Pulendran & Ahmed, 2011).

Finally, because of the generally reduced cell surface protein expression in low SES, we explored the metabolic profile driving these phenotypes by assessing the transcription and translation rates. While lower SES individuals displayed elevated baseline RNA transcription, they exhibited reduced protein translation rates upon activation. This transcriptional-translational uncoupling is consistent with post-transcriptional regulation observed during metabolic stress, chronic inflammation or caloric restriction (Buck et al., 2016; O’Neill et al., 2016). These findings open new avenues for dissecting the molecular basis of immune heterogeneity and linking metabolic state to cellular phenotype, immune functionality and patient outcomes.

Our study has some limitations. The cross-sectional design that we employed precludes definitive causal claims, and while we controlled for age and sex, unmeasured lifestyle variables may account for some variance. The relative low sample size of the discovery cohort was partially addressed by performing validation in independent individuals, validating the key findings. However, there are likely still more subtle differences that were not identified (false-negatives), especially given the large multiple testing correction factor that had to be applied. We also acknowledge that while we measured protein transcription and translation, post-translational modifications that strongly affect protein behavior were not assessed.

In summary, we demonstrated that socioeconomic status is strongly associated with human immune system architecture and cell surface proteome. The use of an unbiased screening approach allowed us to identify hundreds of differentially abundant cell proteins, which could be validated using targeted approaches in additional individuals. It enabled us to identify key altered pathways, related to migration, complement responses, germinal center engagement in T cell functionality. By making the dataset publicly available in an interactive website, we allow for the direct assessment of markers of interest by other researchers. Such an atlas can be of use for predicting potential therapeutic effect of molecules, such as immune checkpoint inhibitors and anti-immune antibodies, in low-SES individuals.

## METHODS

### Study design and participants

We included in this study healthy adults between 18- and 40 years old from Europe (Leiden, the Netherlands) and Sub-Saharan Africa (Senegal).

For Senegalese participants, SES has been defined based on criteria including among others income, education level, housing type, environmental exposure, and healthcare access using questionnaires assessing key SES defining criteria (Evans & Kantrowitz, 2002). In Senegal, low SES participants were recruited in Rufisque (sub-urban, located 19 km away from Dakar) while high SES individuals were selected in Dakar city. Europeans were recruited in Leiden (The Netherlands) and considered as a separate reference group based on their geographical origin and distinct socioeconomic context.

We enrolled 73 participants through the “Geographical Differences of the Immune Response” (GDIR) study In Senegal. This study focussed on changes in the immune profile across Europeans and Senegalese populations between December 2020 and January 2021. The Senegalese participants can be divided in individuals from lower SES backgrounds (Rufisque, n=35) and those from higher SES backgrounds (Dakar city, n=38). Ethical approval was obtained from the National Committee for Ethics and Health Research of Senegal (CNERS, Nr SEN20/56/00000159/MSAS/CNERS/Sec).

European participants were partly recruited through inclusion in the TINO study (n=45 young adults) a prospective cohort study investigating the effects of aging on the immune system, which was conducted at the Leiden University Medical Center (LUMC, Netherlands). The association of immune parameters with covariates as done here was prespecified as an endpoint in this study. Ethical approval was obtained from the Medical Ethical Committee Leiden-Den Haag-Delft (NL77841.058.21). The trial was registered in the Dutch Trial Registry (NL-OMON24946). Additionally, participants were enrolled at the Voluntary Donor Service (LuVDS, LUVDS24.038) at LUMC, where they donated blood anonymously, which we used to develop assays.

All participants participated voluntarily and provided written informed consent according to the Declaration of Helsinki.

### Sample collection and processing

To ensure data quality and comparability across sites, we used standardized study procedures in both sites for sample collection and processing.

From each participant, blood was collected in sodium heparin using standard venipuncture techniques at the Institute of Health Research, Epidemiological Surveillance and Training (IRESSEF, Dakar), (2×10 mL in Dakar, Senegal) and LUMC (8×10 mL in Leiden, the Netherlands). Peripheral blood mononuclear cells (PBMCs) were isolated using density gradient centrifugation with Ficoll. Isolated cells were cryopreserved in a freezing medium containing 100 U/mL penicillin, 100 U/mL streptomycin, 1mM pyruvate, 2mM glutamate, 90% of heat-inactivated Fetal Calf Serum (hi-FCS), and 10% dimethylsulfoxide (DMSO), and cells were then stored in liquid nitrogen. For serological testing, blood was collected in dry or EDTA tube for serum and plasma samples respectively.

PBMCs were isolated within 4 hours following blood collection at all sites using similar protocols. Temperature monitoring systems were used during sample shipment from Senegal to the Netherlands to ensure optimal storage conditions.

### High-throughput spectral flow cytometry profiling using Legendscreen kit

We assessed cell-surface marker expression across phenotypically distinct immune cell subsets from PBMCs in age- and sex-matched individuals, including Europeans (n=7, Leiden) and Senegalese individuals (high SES, n=8; low SES, n=8) (Table S1 and S2). Individuals were selected from clinical studies described above in order to be able to have matching between age and sex between cohorts.

Cryopreserved PBMCs (10 x 10^6 cells) were thawed in pre-warmed Roswell Park Memorial Institute (RPMI) 1640 medium (Gibco™, catalog 11875093) supplemented with 20% hi-FCS at 37°C, washed twice in 5mL of RPMI 20% hi-FCS and transferred to a 96-well V-bottom plate (Nunc™, Catalog 249570). Cells were then pelleted by centrifugation for 5 minutes at 450g and washed twice with 180 μL of Phosphate-Buffered Saline (PBS). After which, cells were resuspended in 100 μL of viability mix containing Fixable Viability Blue Dead (Invitrogen™, catalog L34962, dilution 1:500), Fc receptor binding inhibitor polyclonal antibody (eBioscience™, catalog 16-9161-73, dilution 1:5). We included anti-human CRTH2-BUV805 (dilution 1:50) and anti-human CD27-Super bright 645 (dilution 1:20) antibodies in the viability mix. Cells were incubated for 15 minutes at room temperature, then 80 μL of Fluorescence-Activated Cell Sorting (FACS) buffer (PBS, 0.5% Bovine Serum Albumin [BSA] and 2 mM EDTA) were added, followed by centrifugation. Cells were washed further with 180 μL of FACS buffer.

We performed combinatorial barcoding using anti-human CD45 antibody with four fluorescent dyes, PE-Cy7, Pacific Blue, Alexa Fluor 700, and Brilliant Blue 551 (BioLegend, clone HI30, dilution 1:1600, 1:1600, 1:400, 1:800 respectively). Samples were randomized and analyzed across two consecutive batches using a barcoding scheme that allowed for multiplexing up to 15 samples per batch, along with a Buffy coat reference sample (Sanquin, the Netherlands) included for batch normalization (Fig. S2A). Barcode combinations were added to each sample, incubated on ice for 30 minutes, and washed twice in 180 μL FACS buffer through centrifugation for 5 minutes at 450g. Cells were then resuspended in 180 μL FACS buffer and pooled into a 15 mL Falcon tube pre-filled with 10 mL of FACS buffer and pelleted.

Surface immunophenotyping was performed using a cocktail of 23 monoclonal antibodies (Table S3). Barcoded cell pellets were resuspended in the antibody cocktail and incubated on ice for 30 minutes. After incubation, 80 μL of FACS buffer were added to cells and pelleted for 5 minutes at 450g. Cells were then washed twice with 180 μL of FACS buffer, and pelleted. Next, cells were resuspended in 28.63 mL of FACS in a 50 mL Falcon tube, and transferred to a reagent reservoir for the staining in the LEGENDScreen™ kit.

Following staining with the phenotyping panel, we performed the preparation and the staining procedure for the LEGENDScreen™ kit according to the manufacturer’s recommendations. Briefly, 75 μL of cell suspension was dispensed into each well of the LEGENDScreen™ kit (Biolegend, catalog 700011, Lot № 380207). The kit is provided in 4 x 96-well V-bottom plates containing 354 reconstituted PE-conjugated antibodies, as well as 10 isotype controls (complete list in Table S4). We skipped wells that contain markers already included in the phenotyping panel, leading to 331 markers that were analyzed. Fixed cells were resuspended in 150 μL of FACS buffer and transferred into 1.5 mL FACS tubes. Acquisition was performed on the Cytek® Aurora 5 laser cytometer (Cytek Biosciences; acquisition software, SpectroFlo, v3.0) at the LUMC flow cytometry core facility.

### Validation of SES-related immune markers

We validated the key SES-related markers from the discovery cohort in an independent cohort (Table S5 and S6) using three panels targeting key markers identified in B cells (Table S7), myeloid cells (Table S8), and T cells (Table S9), respectively. The validation cohort included age- and sex-matched participants (n=10 per group: European, high SES Senegalese, low SES Senegalese) processed using identical protocols.

Cryopreserved PBMCs (10 x 10^6 cells) were thawed in pre-warmed RMPI 20% hi-FCS at 37°C. PBMCs were stained with all three targeted panels in parallel where for each donor 5 x 10^5 cells were plated in 96-well V-bottom plate. Cells were washed twice with 180 μL of PBS and pelleted through centrifugation for 5 minutes at 450g. Next, cells were resuspended in 50 μL of Fixable Viability Blue Dead (Invitrogen™, catalog L34962) with Fc-R blocker (eBioscience™, catalog 16-9161-73), and incubated for 15 minutes at room temperature, then subsequently 50 μL of 2X concentrated antibody cocktail from each panel was added to cells, and incubated on ice for 30 minutes. After incubation, 80 μL of FACS buffer were added to the cells, followed by a 5 minutes centrifugation at 450g. Cells were washed twice with 180 μL of FACS, pelleted by centrifugation, and fixed with 4% Ultra-Pure Formaldehyde (Thermo Scientific™, catalog 28908) for 10 minutes at room temperature. After fixation, cells were pelleted through centrifugation for 5 minutes at 800g. Cells were then washed with 180 μL of FACS buffer, and resuspended in 150 μL of FACS and transferred to 1.5 mL FACS tubes for acquisition on Cytek® Aurora 5 laser cytometer.

#### Flow cytometry data analysis

Flow cytometry data were unmixed and compensated using Cytek SpectroFlo software (v3.0). Immune cell subsets of interest were defined by manual gating after excluding the singlets and dead cell using FlowJo (v10.10.0) and the cloud-based platform OMIQ (Dotmatics, 2025), the detailed gating strategy is shown in Figure S1. Next, we conducted the de-barcoding of the samples using custom scripts written in R v.4.4.3 (R Core Team, 2025), utilizing the RStudio interface v.2024.12.1+563(Posit team, 2025) with R flowCore package v1.11.20, where we defined a cutoff for each barcode per cell population separately. Cells were assigned to a individual sample based on the positivity for each of the 4 barcodes according to previously assigned barcode combinatorial (Fig. S2B-C). One lower SES participant (1/8) was excluded at this point due to inadequate cellular yield (n < 5000 cells per well). We normalized PE signal intensity for all markers by subtracting the background signal from each corresponding isotype control. For each immune subset and sample, we exported the median fluorescence intensity (MFI) and the 90^th^ percentile. Only markers with an MFI above a threshold of 1000 were included the statistical analysis. High-dimensional clustering analysis was performed using *FlowSOM* algorithm (Van Gassen et al., 2015). Dimensionality reduction was performed with R umap package v0.2.10.0 using Uniform Manifold Approximation and Projection (UMAP) for visualization (McInnes & Healy, 2018).

### Serum complement activation analysis

Complement activity of sera was measured using in house enzyme-linked immunosorbent assay (ELISA) (van den Beukel et al., 2025). To assess complement activation in each pathway, 96-well plates were coated overnight at 4°C with intravenous immunoglobulin (IVIG, 10 μg/mL) for the classical pathway, acetylated bovine serum albumin (BSA, 10 μg/mL) for the lectin pathway, or lipopolysaccharide (LPS, 10 μg/mL) for the alternative pathway. Following coating, plates were blocked with PBS containing 1% bovine serum albumin for 1 hour at room temperature. Serum samples were serially diluted in RPMI supplemented with MgEGTA for the alternative pathway, or in RPMI for the classical and lectin pathways, and added to the coated plates. Plates were incubated for 1 hour at 37°C to allow complement activation and C5b-9 deposition. As a marker of terminal complement activation, plate-bound C5b-9 complexes were detected using mouse anti-human C5b-9 monoclonal antibody (Dako, M0777) and horseradish peroxidase-conjugated (HRP) goat anti-mouse immunoglobulin (Dako, P0447). The enzymatic reaction was developed using ABTS (Merck, A1888-5G) substrate with H_2_O_2_ (Merck, 1072090250), and absorbance was measured at 415 nm using a microplate reader (BIO-RAD iMark). Normal human serum (NHS) was used as the positive control, and heat-inactivated serum as the negative control. Data were normalized to NHS values at optimal dilutions (10% for the alternative, 0.3% for the classical, and 3% for the lectin). The normalized optical density values representing relative C5b9 binding and reflecting complement activation in each pathway were compared across the group.

### Metabolic labeling of newly synthesized proteins and RNA using a bioorthogonal analog

We measured protein translation rates using bioorthogonal non-canonical amino acid (ncAA) labeling with β-ethynylserine (βES, provided by Kimberly Bonger, University of Leiden) in PBMCs from an additional cohort (n = 10 per group) including Senegalese stratified by SES along with European participants recruited at the LUVDs (Table S10). βES was used as described in previously published methods (Ignacio et al., 2023; Vrieling et al., 2024).

PBMCs were cultured in RPMI medium supplemented with 10% hi-FCS for 1 hour at 37°C and 5% CO_2_. Cells were then treated with 500 µM βES for 1 hour under the same conditions. For stimulation, PBMCs were incubated overnight with a mixture containing 1 µM CPG-ODN2006 (Invitrogen, catalog tlrl-2006), 100 ng/mL LPS (InvivoGen, catalog tlrl-3pelps), 1 µg/mL purified mouse anti-human CD3 (BD Pharmagen, catalog 555336), and 10 µg/mL anti-human CD28 (BD Pharmagen, catalog 555725) in

RPMI medium with 10% hi-FCS, followed by βES treatment as above. As a control for βES incorporation during protein synthesis, samples were pre-treated with 100 µM Cycloheximide (CHX, Tocris Biosciences, Cat#0970), a protein synthesis inhibitor.

After incubation, cells were pelleted by centrifugation for 5 minutes at 450g and washed twice with 180 μL of PBS. Cell pellets were incubated for 15 min at room temperature with 50 μL of cocktail mix containing Zombie Violet™ Fixable Viability Dye (Biolegend, catalog 423113, dilution 1:2000), Fc-R blocker and monoclonal antibodies for surface immunophenotyping and (Table S11). Upon incubation, 130 μL of FACS were added to cells, followed by centrifugation for 5 minutes at 450g and washed with 180 μL of PBS. Subsequently cells were fixed with 180 μL of 2% Ultra-Pure Formaldehyde, followed by centrifugation for 5 minutes at 800g, then cells were washed with 180 μL of PBS.

Fixed cells were permeabilized by incubation in with 180 μL of 1X eBioscience^TM^ Permeabilization Buffer (Invitrogen, catalog 00-8333-56) for 15 minutes at room temperature. Fluorescent detection was conducted via click chemistry for 30 minutes at room temperature. Cells were resuspend in 50 μL of click reaction mix containing 0.5 mM copper sulfate (CuSO_4_, Bonger Lab), 2 mM Tris-hydroxypropyltriazolylmethylamine (THPTA, Bonger Lab), 10 mM sodium ascorbate (NaAsc, Bonger Lab), and 0.25 µM AZDye 647 Azide Plus (Vector Laboratories, catalog CCT-1482-1) in PBS. After incubation, 130 μL of FACS were added to cells, followed by centrifugation for 5 minutes at 800g and washed with 180 μL of PBS. We performed intracellular staining of CD3 Spark Blue 574 (Biolegend, catalog 300487, dilution 1:50 in 1X Perm buffer) in the presence of Fc-R blocker and BD Stain Plus (BD, catalog 566385, dilution 1:10).

In parallel, we applied the same approach to label newly transcribed RNA by measuring 5-ethynyl uridine (5’EU, Tocris Biosciences, catalog 7206) incorporation, a uridine analog that serves as a proxy for RNA transcription. To confirm that 5’EU labels cellular RNA, cells were incubated with 20 µM Actinomycin D (Act D, Tocris Biosciences, Cat#1229) to evaluate the effect of RNA polymerase inhibition on 5’EU labeling. Following resting and stimulation, PBMC were treated with 500 µM of 5’EU in RPMI 10% FCS for 1h at 37°C and 5% CO_2_ to assess its incorporation into newly transcribed RNA. Fluorescent detection was performed via click chemistry using 0.25 µM AZDye 647 Azide Plus (Jao & Salic, 2008; Van’t Sant et al., 2021).

### Statistical analysis

Linear models were fitted to log-transformed marker expression values within each immune cell subset using the R limma package (R version 4.2.2, RStudio version 2024.04.2+764). To account for varying precision across samples, we weighted the model by immune subset counts, reflecting the greater reliability of more abundant immune subsets. We tested for monotonic trends across groups (European, High SES, Low SES) by modelling group as an ordinal variable, under the assumption that SES behaves as a gradient and Europeans reflect the highest SES group. Such a design is powerful for detection of monotonic trends but may miss non-monotonic patterns. We applied this statistical model to compare immune cell subset frequencies as well as for assessing changes in the metabolic rates between groups. The models included batch as a covariate, with additional batch correction applied when needed using CytoNorm, an Omiq.ai normalization algorithm for cytometry data platform (Van Gassen et al., 2020). P-values were subsequently adjusted for multiple testing using the Benjamini-Hochberg method across all tested markers and populations (*P* < 0.05). For complement activity assay, we performed statistical analysis using the Kruskal-Wallis test for group comparisons, with *P* < 0.05 corrected for multiple comparisons considered significant. Post-hoc pairwise comparisons were conducted using the Wilcoxon test.

## Acknowledgements

The authors thank all clinical and research staff at UCAD and IRESSEF (Dakar, Senegal) and the LUMC (Leiden, the Netherlands) for their essential contribution to participant recruitment. We also thank the LUMC Core facility for providing spectral flow cytometry services. Finally, our sincere thanks go to all volunteers who participated in this study, this work would not been possible without them.

## Author Contributions

MM, MY and SPJ designed the study.

MC, ID, MG and MM organised the recruitment and collected samples from Senegalese participants. MC, ID and MG processed the samples from Senegal.

WH, GG, SM, SPJ organised the recruitment and collected samples from the European participants. MC, WH, SS and AK conducted spectral flow cytometry experiments.

MC, HW and LAT carried out / interpreted experiments on complement activation. KB provided BES-HCL analog for bioorthogonal labelling of nascent proteins.

MC performed the experiments for the click chemistry labelling of nascent metabolites. MC and SPJ analysed the data.

MC and YS developed the Shiny interactive dashboard.

MC wrote the first draft of the manuscript, SPJ, MM and MY contributed to subsequent versions. SPJ, MC, and MM verified all the data reported in this study.

MM, SPJ, and MY oversaw the project.

All authors reviewed, and approved the manuscript.

These authors jointly supervised this work: M.M., S.P.J., and M.Y.

## Data availability and shiny app (SES - Atlas)

All data needed to evaluate the conclusions in the paper are present in the paper or the Supplementary Materials. Processed flow cytometry datasets are deposited in the Zenodo repository, at https://doi.org/10.5281/zenodo.17063250. Data are deposited under restricted access to comply with ethical and privacy regulations. Researchers will submit their research idea or protocol to the corresponding authors to ensure it aligns with approved protocols and consent forms. Once approved, data access will be granted without any fees to be charged.

We deployed an interactive web application built with custom scripts using the R Shiny package (version 1.10.0) to explore immunological profiles across SES groups, accessible at https://cismar.shinyapps.io/LegendscreenShinyApp/. This web tool allows users to visualize marker expression patterns across different immune cell subsets and compare expression levels between SES groups. All code used to build the Shiny application including all packages, functions, and key parameters used are publicly available at https://doi.org/10.5281/zenodo.18063314.

---

**Fig. S1.**
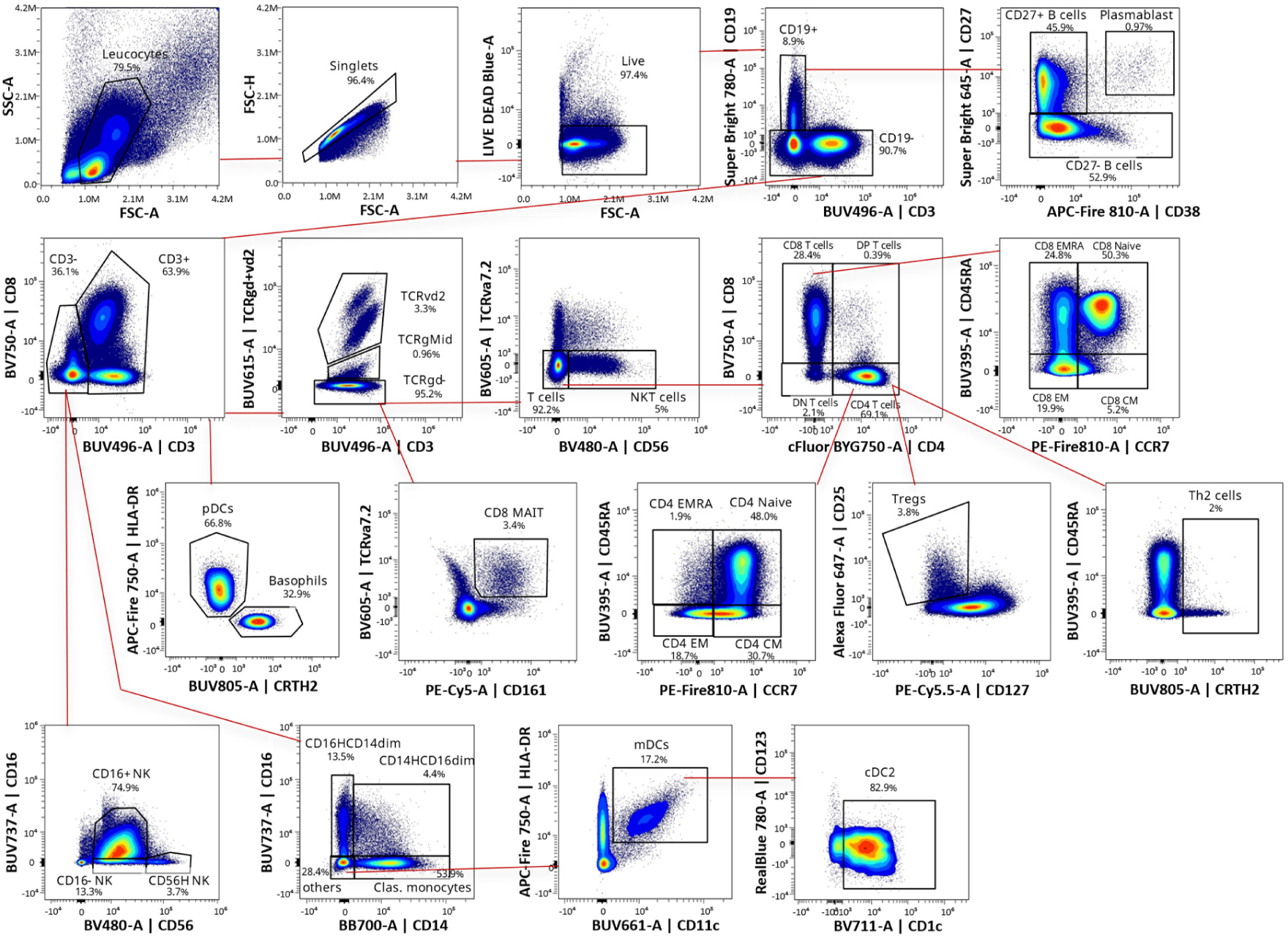
Flow cytometry gating strategy of peripheral immune cell subsets. Overview of manual gating analysis used to identify 38 immune cell subsets based on established surface marker expression profiles from the phenotyping panel provide in Table Supplementary 1. Following doublets (FSC-H vs FSC-A) and dead cells exclusion (LIVE DEAD Blue vs FSC-A), major cell types were identified and further differentiated into unique subsets. This included B cells subsets (CD27Pos B cells : CD27^+^ CD38^-^, CD27Neg B cells : CD27^-^ CD38^+/-^, and Plasmablast : CD38^high^ CD27^+^) as well as CD4+ T cells (Tregs : CD25^+^ CD127^-^, Th2 cells : CRTH2^+^, and CD161^+^ CM/EM) and CD8^+^ T cells with theirs memory subsets (Naïve : CD45RA^+^ CCR7^+^, EM : CD45RA^-^ CCR7^-^, CM : CD45RA^-^ CCR7^+^, EMRA : CD45RA^+^ CCR7^-^). Unconventional T cells (DPT cells : CD3^+^ CD4^+^ CD8^+^, DNT cells : CD3^+^ CD4^-^ CD8^-^, γδ T cells : TCRgd/vd2^+^, MAIT cells : TCRva7.2^+^ CD161^+^, and invariant NKT cells : CD3^+^ CD56^+^ CD4/CD8^+^) were also defined as were NK cell subsets (CD56H NK cells : CD56^bright^ CD16^-^, CD16Pos NK cells : CD56^+^CD16^+^, and CD16Neg NK cells : CD56^dim^ CD16^-^), Monocytes (Classical monocytes : CD14^+^ CD16^dim^, Interm. monocytes : CD14^+^ CD16^+^ and Non clas. monocytes : CD14^dim^ CD16^high^), Dendritic cells (mDCs : HLA-DR^+^ CD11c^+^, cDC2 : HLA-DR^+^ CD11c^+^ CD1c^+^ and pDCs : HLA-DR^+^ CD123^+^ CD11c^-^) and Basophils (CD123^+^ CRTH2^+^ HLA-DR^-^).

**Fig. S2.**
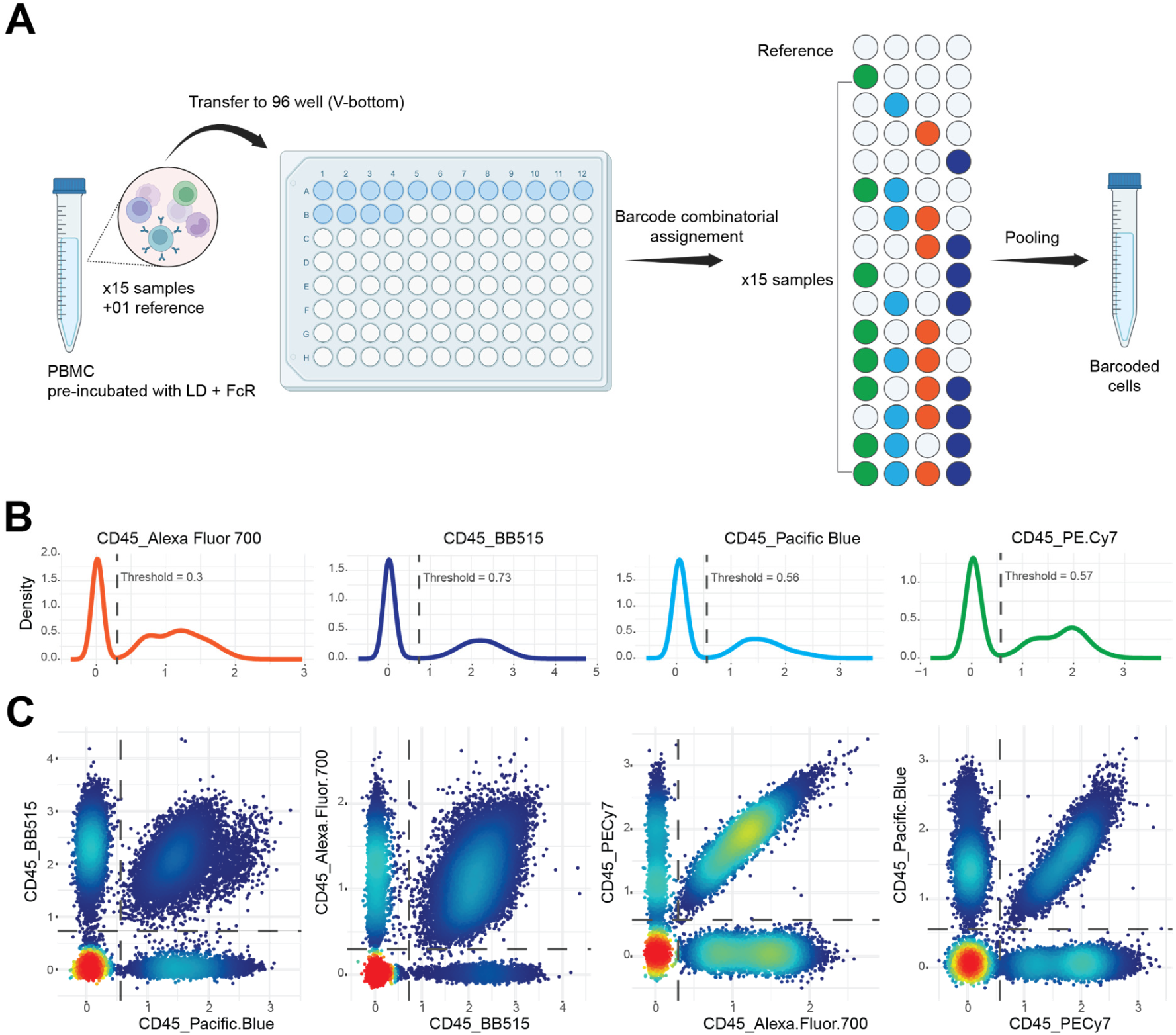
Combinatorial barcoding and demultiplexing of study participants based on CD45 expression. **A**) Workflow overview of sample barcoding using anti-human CD45 antibody coupled with four unique fluorochromes enabling combinatorial labeling of participant samples. **B**) Histograms displaying CD45 signal intensity for each barcode. Dashed lines indicate the fluorescence thresholds used to delineate positive and negative populations for each unique fluorochrome. **C**) Debarcoding of study cohort participants based on initial barcode combinations using density gating.

**Fig. S3.**
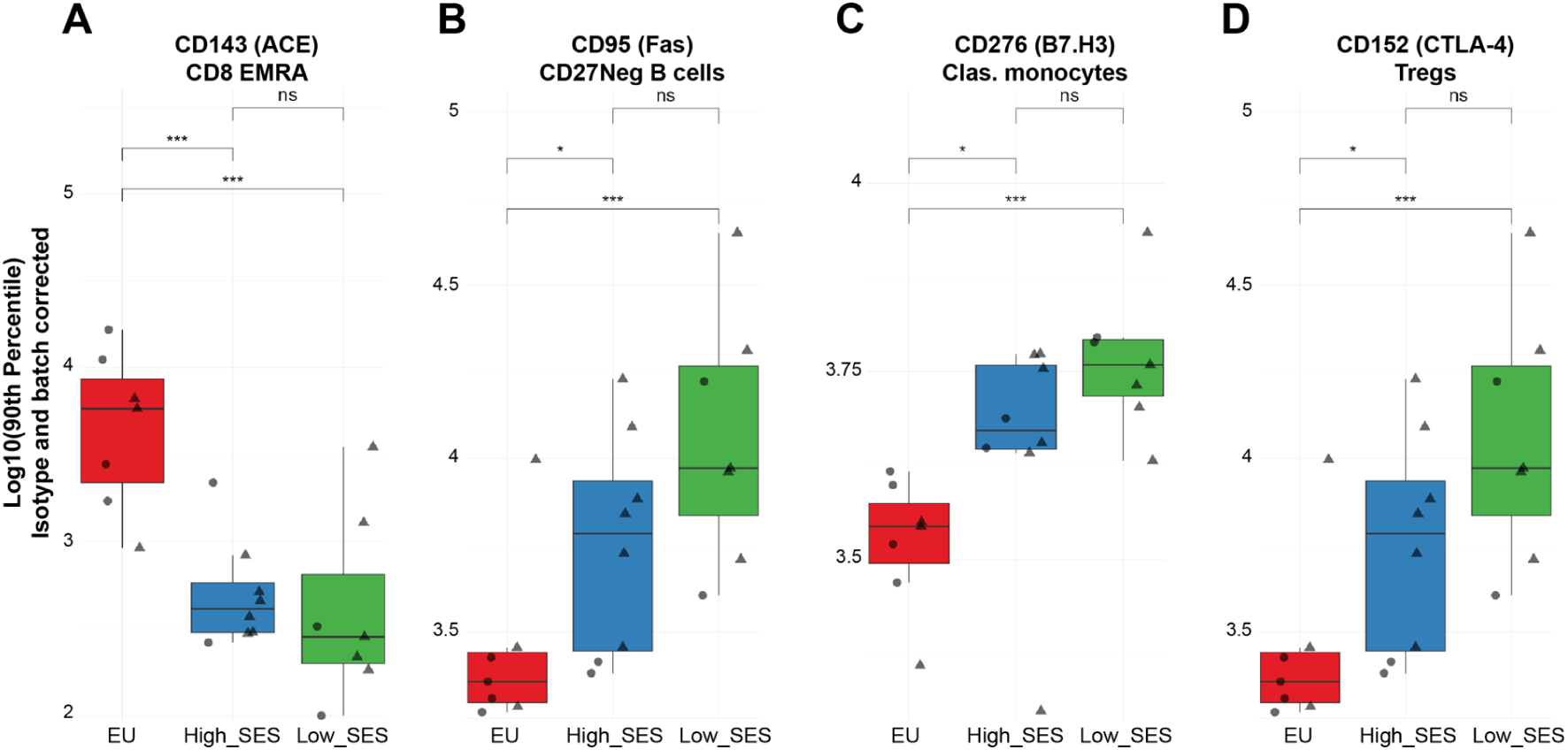
90th percentile expression analysis reveals exclusive SES-related markers. Boxplots showing the 90^th^ percentile of expression of surface proteins with significant changes along SES gradients that were not captured by MFI analysis. **A**-**D**) Example of the angiotensin converting enzyme, CD143 (on CD8+ EMRA), the Fas receptor (on CD27Neg B cells), the immune checkpoint proteins B7.H3 as well as CTLA-4 are shown.

**Fig. S4.**
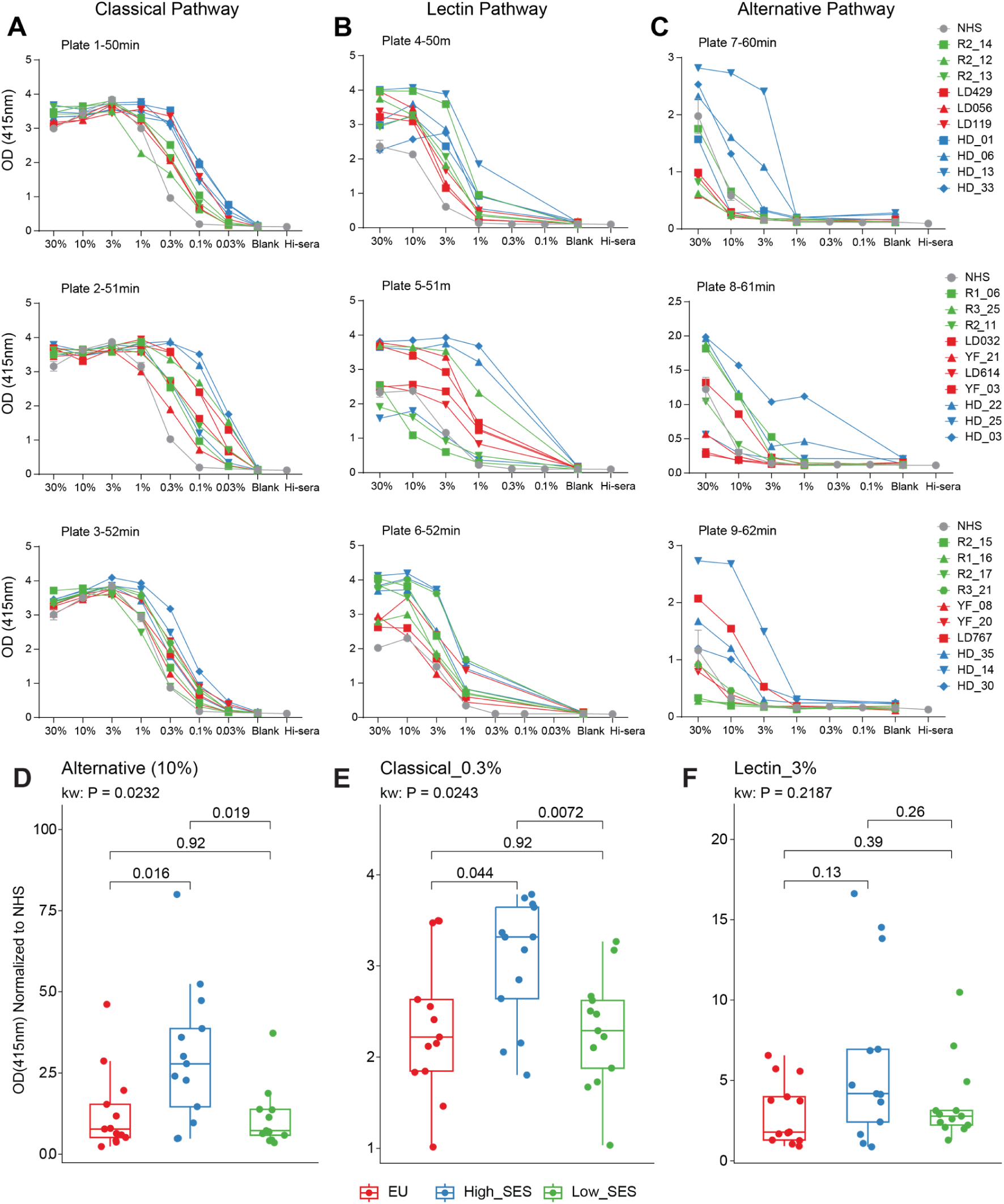
Complement activity is enhanced in higher-SES Senegalese populations. Complement activation was assessed using pathway-specific plate-bound assays. (**A**-**C**) Representative titration curves showing C5b-9 detection in serum samples for (**A**) classical pathway (Immunoglobulin G-coated wells), (**B**) lectin pathway (acetylated bovine serum albumin-coated wells), and (**C**) alternative pathway (Lipopolysaccharide-coated wells). Boxplots showing complement activity normalized to normal human serum (NHS) across SES groups for (**D**) alternative pathway at 10% serum dilution, (**E**) classical pathway at 0.3% serum dilution and (**F**) lectin pathway at 3% serum dilution. Statistical significance was determined using the Kruskal-Wallis test (*P* < 0.05) followed with pairwise comparison using Wilcoxon test, corrected for multiple testing.

**Fig. S5.**
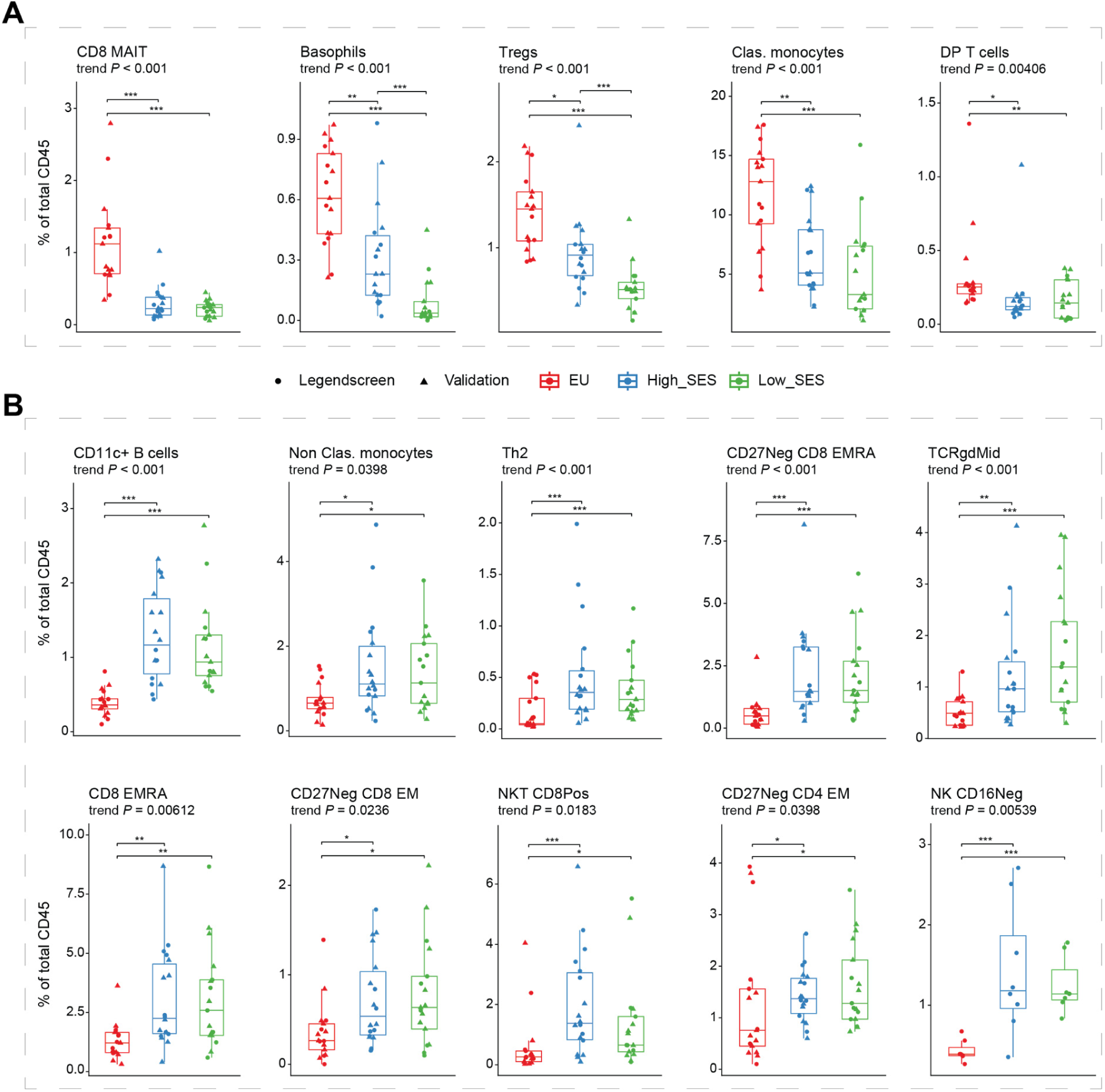
Immune subset frequencies correlate with socioeconomic status. Boxplots of frequencies of manually gated immune subsets showing SES-dependent changes across phenotyping and targeted panels (trend analysis FDR < 0.05). **A**) Immune subsets significantly associated with higher SES. **B**) Differentially abundant immune subsets related to lower SES.

**Fig. S6.**
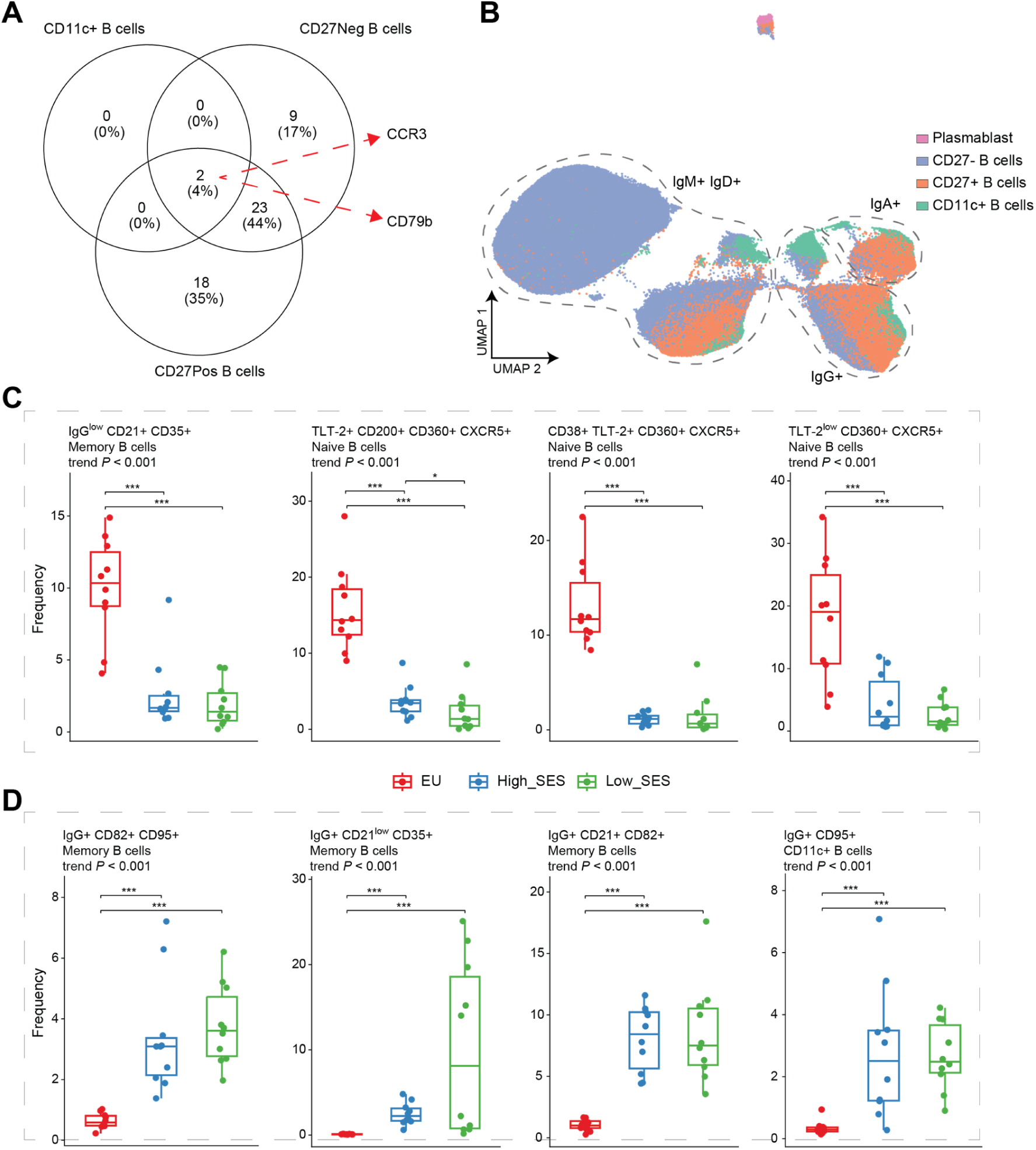
Unsupervised clustering analysis reveals SES-associated B cell subsets. **A)** Venn diagram showing the number of overlapping markers between B cell subsets. **B)** UMAP dimensionality reduction plots displaying the distribution of immune cell subsets of interest in the validation cohort. Subsets were identified through manual gating and projected onto two-dimensional UMAP space. Distributions are shown for B cell subsets. Boxplots showing frequencies of differentially abundant B cell clusters between the groups (trend analysis FDR < 0.05). **C**) B cell clusters highly enriched in individuals from higher SES backgrounds. **D**) Differentially abundant B cell clusters related to lower SES.

**Fig. S7.**
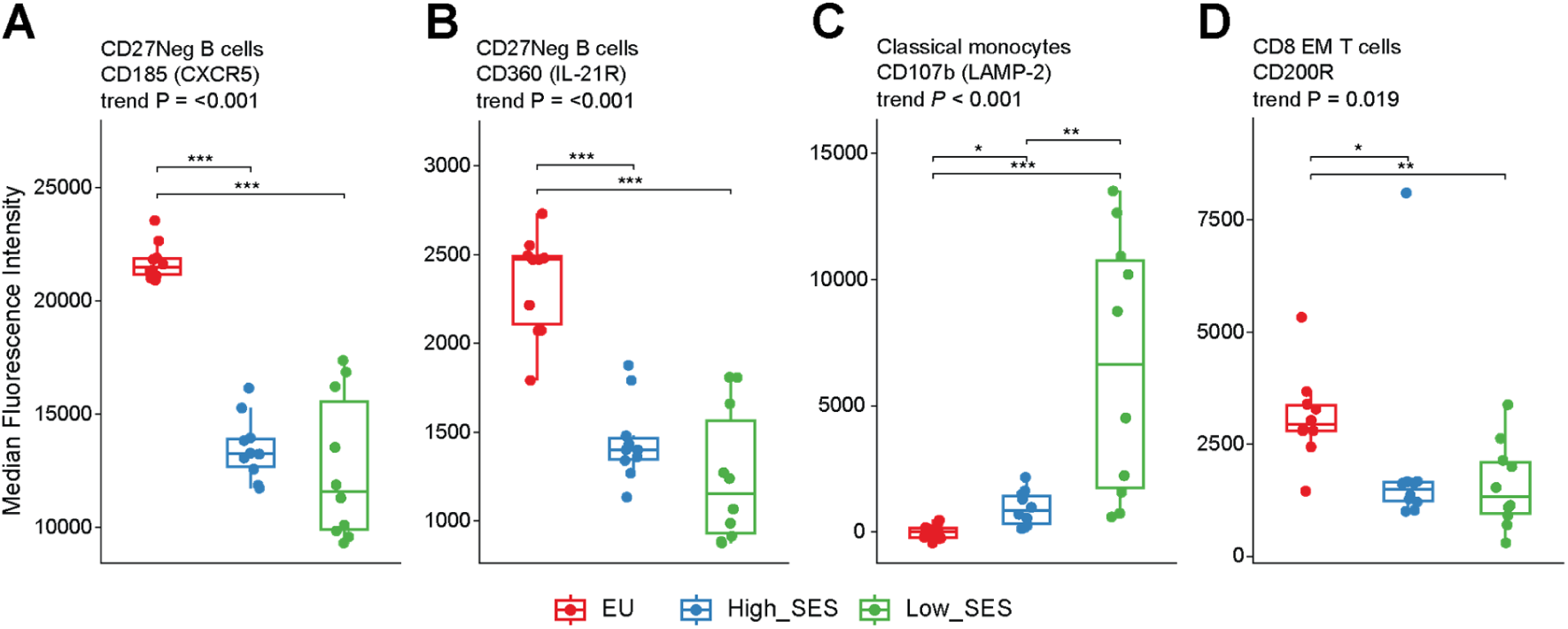
Validation cohort confirms LEGENDScreen findings on SES-related surface proteins. Boxplots showing the median fluorescence intensity of differentially expressed cell surface protein from targeted validation panels exhibiting SES-dependent changes between the groups (trend analysis FDR < 0.05) consistent with the legendscreen findings. **A-D**) Example of the chemokine receptor CXCR5 and IL-21R (on CD27Neg B cells) are shown together with LAMP-2 (on Clas. monocytes) and the immune checkpoint CD200R (on CD8 EM T cells).

**Fig. S8.**
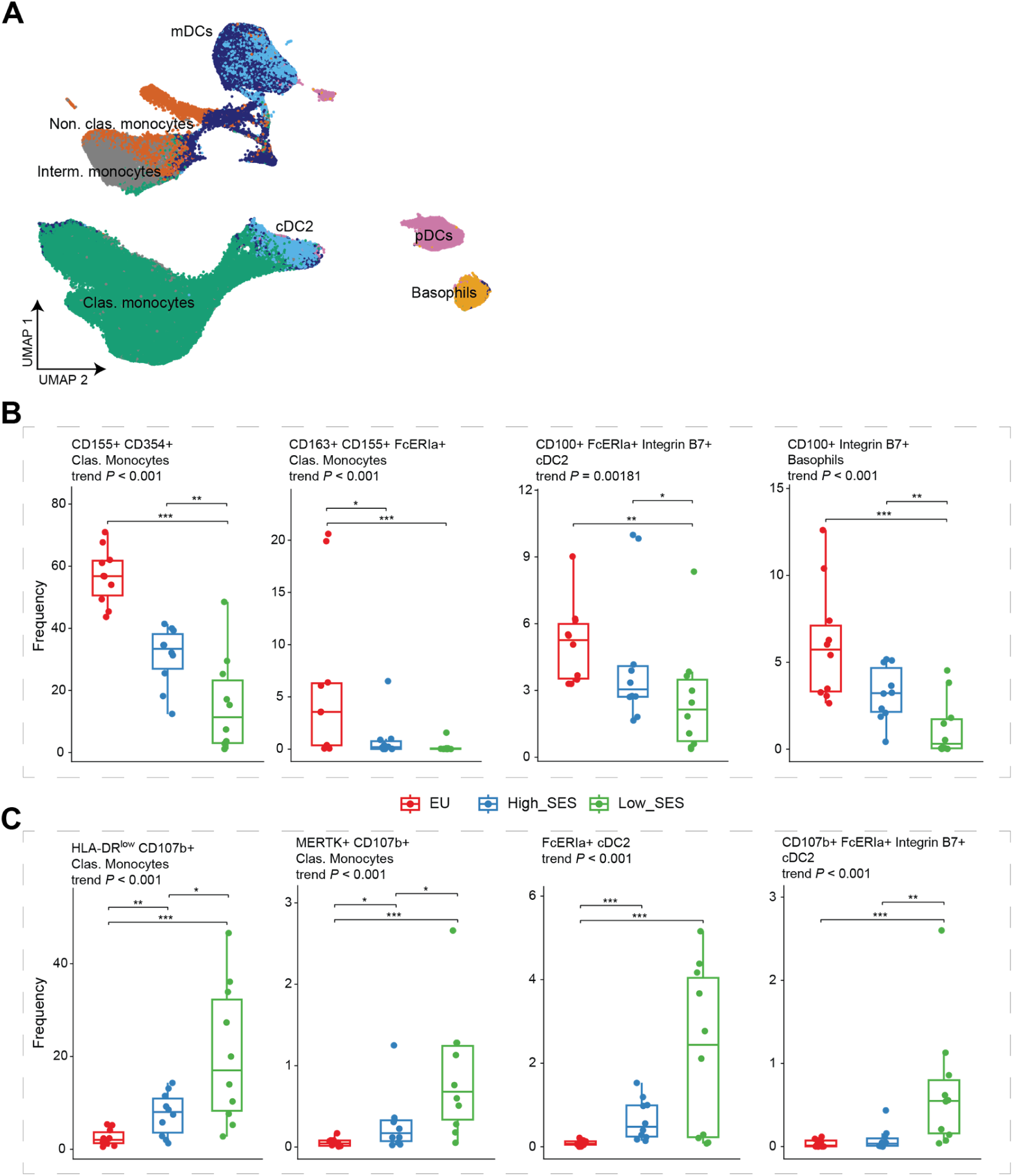
Targeted myeloid validation panel identifies clusters differentiating between the SES gradients. **A**) UMAP dimensionality reduction plot displaying the distribution of myeloid cell subsets of the validation cohort. Subsets were identified via manual gating and projected onto two-dimensional UMAP space. Distributions of myeloid cell subsets are shown. Boxplots showing frequencies of differentially abundant myeloid cell clusters between the groups (trend analysis FDR < 0.05). **B**) Clusters frequencies associated with higher SES. **C**) Clusters abundance related to lower SES.

**Fig. S9.**
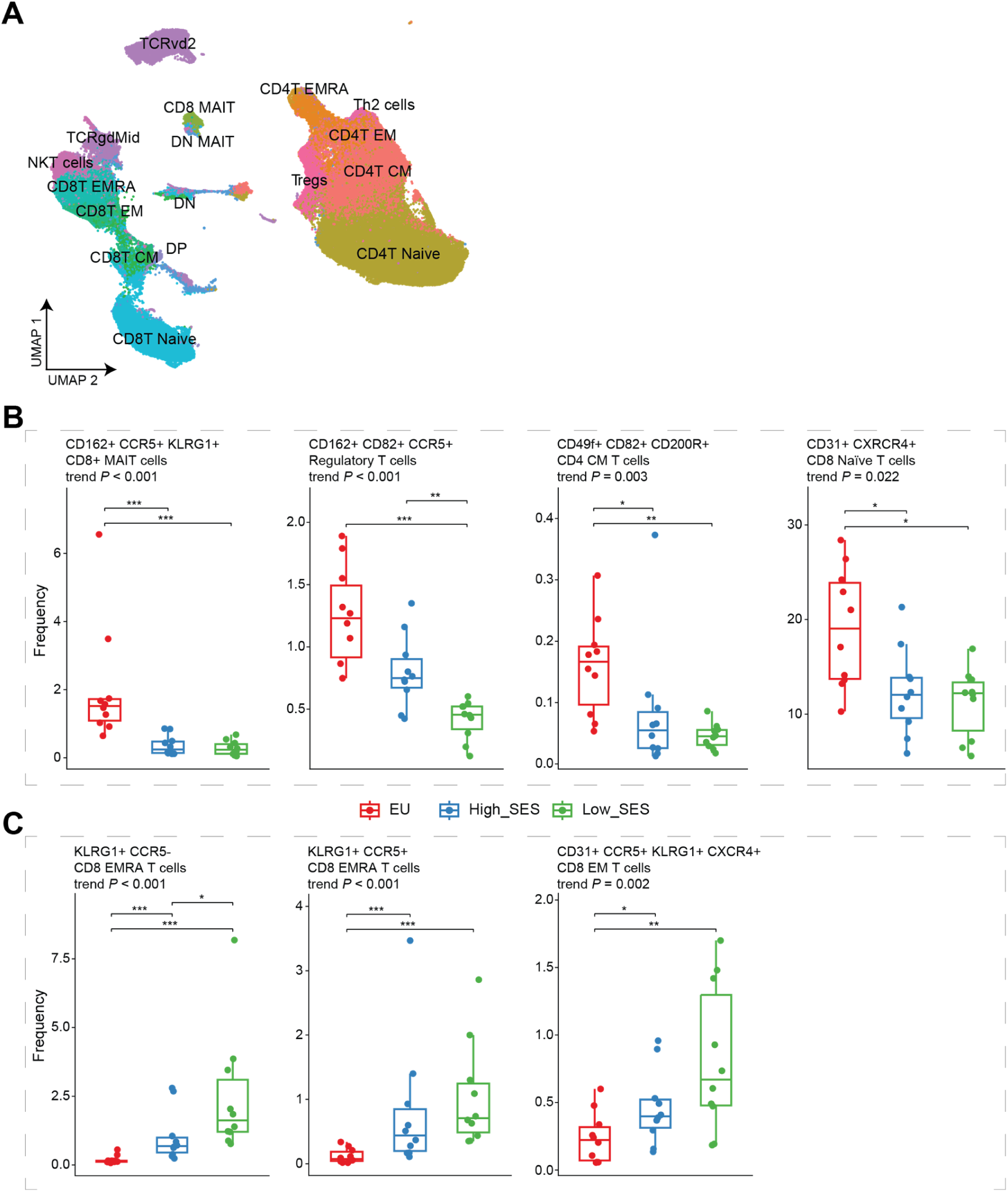
Targeted T cell validation panel identifies clusters differentiating between the SES gradients. **A**) UMAP dimensionality reduction plot showing the distribution of T cell subsets of the validation cohort. T cell subsets were identified via manual gating and projected onto two-dimensional UMAP space. Boxplots showing frequencies of differentially abundant T cell clusters between the groups (trend analysis FDR < 0.05). **B**) B clusters highly enriched in individuals from higher SES backgrounds. **C**) Differentially abundant T cell clusters related to lower SES.

**Fig. S10.**
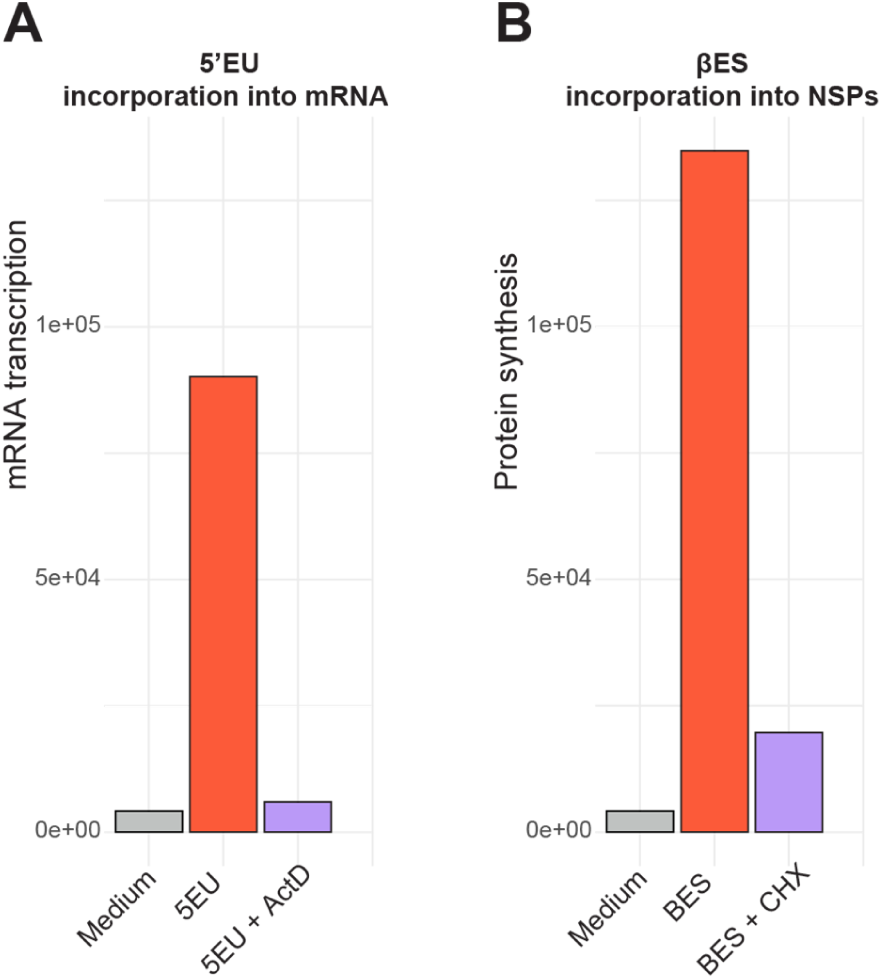
Targeted inhibition of transcription and translation validates probes specificity. To validate the specificity of the metabolic probes, cells were incubated with medium only or 5’EU with and without 20 µM of actinomycin D (Act D) as a control for RNA transcription (**A**). Similarly, as control for βES incorporation, cells were incubated in the presence or absence of 100 µM of cycloheximide (**B**).

**Table S1.** Demographics of discovery cohorts.

| Characteristic | Residence |  |  |
| --- | --- | --- | --- |
|  | EU<br>N = 7<sup>1</sup> | High SES<br>N = 8<sup>1</sup> | Low SES<br>N = 8<sup>1</sup> |
| **Age, years** | 29.0 (24.0 - 35.0) | 29.0 (25.0 - 33.0) | 29.0 (23.0 - 35.0) |
| **Sex, female** | 3 (43%) | 2 (25%) | 4 (50%) |
| **Height, cm** | NR | 1.78 (0.10) | 1.74 (0.08) |
| *Missing* | 7 | 0 | 4 |
| **Weight, kg** | NR | 75 (7) | 69 (15) |
| *Missing* | 7 | 0 | 0 |
| **BMI, kg/m<sup>2</sup>** | NR | 23.92 (4.24) | 24.41 (1.82) |
| *Missing* | 7 | 0 | 4 |
| **Systolic blood pressure, mmHg** | NR | 120 (110, 130) | 120 (111, 124) |
| *Missing* | 7 | 0 | 0 |
| **Diastolic blood pressure, mmHg** | NR | 80 (80, 85) | 69 (65, 83) |
| *Missing* | 7 | 0 | 0 |
| **Current or past tobacco use** | 0 (0%) | 4 (50%) | 0 (0%) |
| **Passive smoking** | 0 (0%) | 8 (100%) | 3 (38%) |

<sup>1</sup>Median (Min - Max); n (%); Mean (SD); Median (Q1, Q3)

**Table S2.** Socio-economic status indicators of Senegalese Legendscreen discovery cohorts.

| **Characteristic** | **Residence** |  | **p-value<sup>2</sup>** |
| --- | --- | --- | --- |
|  | **High SES**<br>**N = 8<sup>1</sup>** | **Low SES**<br>**N = 8<sup>1</sup>** |  |
| Lived in current residence ≥5 years | 8 (100%) | 8 (100%) |  |
| Years spent in current residence category, normalized by age | 0.93 (0.68) | 1.00 (1.00) | 0.2 |
| Mean number of people living in the same house | 5.1 (2.0) | 15.9 (2.9) | <0.001 |
| **Education level** |  |  | 0.2 |
| Elementary | 0 (0%) | 1 (13%) |  |
| Secondary | 3 (38%) | 4 (50%) |  |
| Uninstructed | 1 (13%) | 3 (38%) |  |
| University | 4 (50%) | 0 (0%) |  |
| **Housing type** |  |  | 0.001 |
| Appartement | 8 (100%) | 1 (13%) |  |
| House with concessions | 0 (0%) | 7 (88%) |  |
| **Floor coverings** |  |  | <0.001 |
| No | 0 (0%) | 6 (75%) |  |
| Partial | 0 (0%) | 2 (25%) |  |
| Yes | 8 (100%) | 0 (0%) |  |
| **Wall coverings** |  |  | 0.001 |
| No | 0 (0%) | 4 (50%) |  |
| Partial | 0 (0%) | 3 (38%) |  |
| Yes | 8 (100%) | 1 (13%) |  |
| **Frequency of Incomes** |  |  | 0.026 |
| irregular | 3 (38%) | 7 (100%) |  |
| regular | 5 (63%) | 0 (0%) |  |
| Missing | 0 | 1 |  |
| **Contact with animals** | 1 (13%) | 3 (60%) | 0.2 |

| Characteristic | Residence |  | p-value<sup>2</sup> |
| --- | --- | --- | --- |
|  | High SES<br>N = 8<sup>1</sup> | Low SES<br>N = 8<sup>1</sup> |  |
| Missing | 0 | 3 |  |
| **Attitude when sick** |  |  | 0.018 |
| automedication | 1 (13%) | 2 (25%) |  |
| see a doctor | 7 (88%) | 2 (25%) |  |
| wait to feel better | 0 (0%) | 4 (50%) |  |

<sup>1</sup>n (%); Mean (Min); Mean (SD)
<sup>2</sup>NA; Wilcoxon rank sum test; Fisher's exact test

**Table S3.**
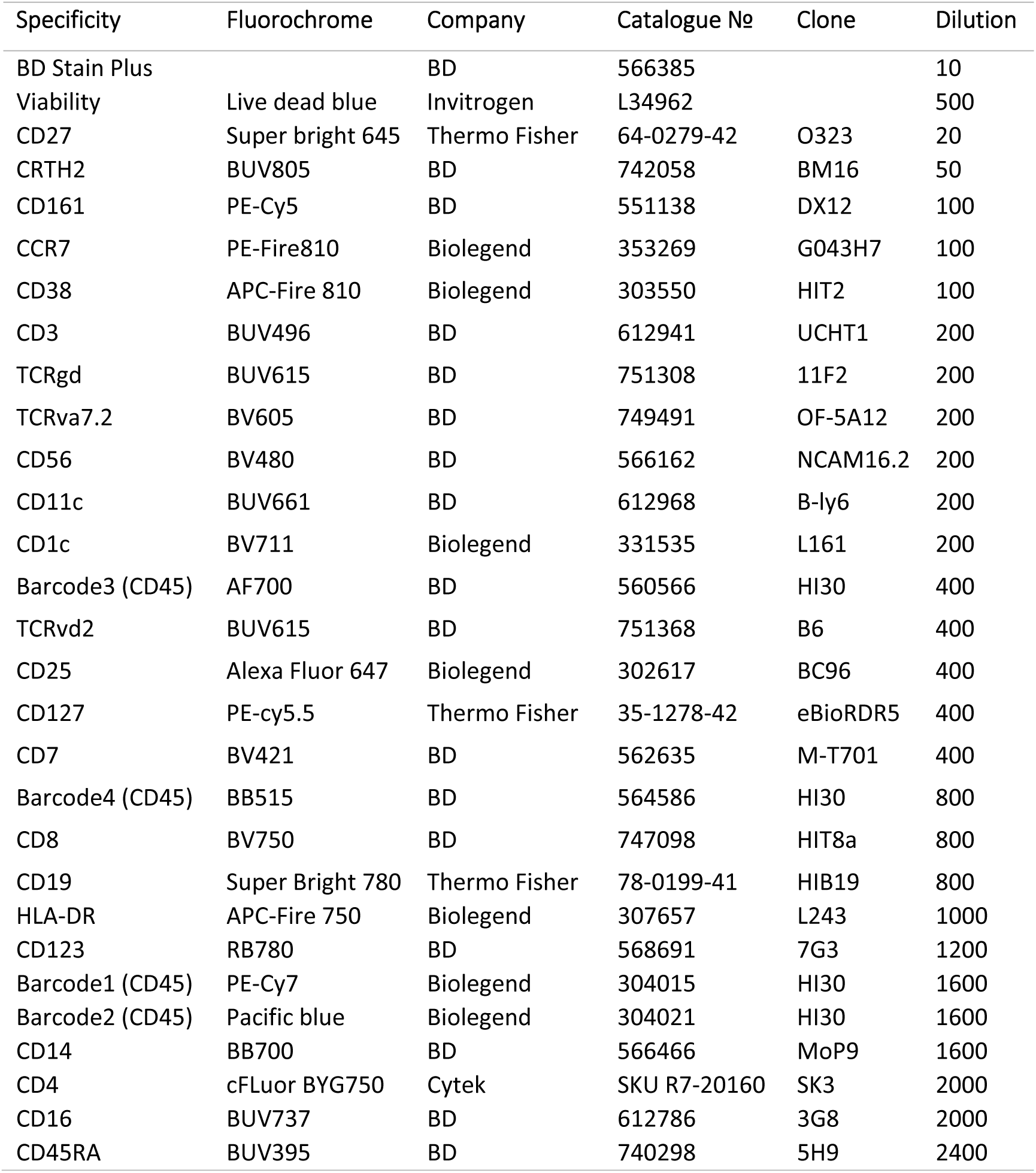
Spectral flow cytometry panel for phenotyping of peripheral immune cells.

**Table S4.** Human PE-conjugated LEGENDscreen antibodies kit (see excel sheet)

**Table S5.** Demography of flow cytometric validation cohort.

| **Characteristic** | **Residence** |  |  |
| --- | --- | --- | --- |
|  | **EU**<br>N = 10<sup>1</sup> | **High SES**<br>N = 10<sup>1</sup> | **Low SES**<br>N = 10<sup>1</sup> |
| **Age, years** | 26.0 (20.0 - 37.0) | 26.0 (23.0 - 37.0) | 26.0 (21.0 - 37.0) |
| **Sex, female** | 6 (60%) | 6 (60%) | 6 (60%) |
| **Height, cm** | NR | 1.74 (0.10) | 1.76 (0.07) |
| *Missing* | 10 | 1 | 5 |
| **Weight, kg** | NR | 75 (7) | 69 (12) |
| *Missing* | 10 | 2 | 0 |
| **BMI, kg/m<sup>2</sup>** | NR | 24.5 (3.0) | 22.2 (4.2) |
| *Missing* | 10 | 2 | 5 |
| **Systolic blood pressure, mmHg** | NR | 110 (110, 120) | 119 (117, 122) |
| *Missing* | 10 | 1 | 0 |
| **Diastolic blood pressure, mmHg** | NR | 80 (80, 80) | 78 (73, 82) |
| *Missing* | 10 | 1 | 0 |
| **Current or past tobacco use** | 0 (0%) | 2 (20%) | 0 (0%) |
| **Passive smoking** | 0 (0%) | 8 (80%) | 6 (60%) |

<sup>1</sup>Median (Min - Max); n (%); Mean (SD); Median (Q1, Q3)

**Table S6.** Socio-economic status indicators of Senegalese validation cohorts.

| **Characteristic** | **Residence** |  | **p-value<sup>2</sup>** |
| --- | --- | --- | --- |
|  | **High SES**<br>**N = 10<sup>1</sup>** | **Low SES**<br>**N = 10<sup>1</sup>** |  |
| Lived in current residence ≥5 years | 7 (70%) | 10 (100%) | 0.2 |
| Years spent in current residence category, normalized by age | 0.55 (0.08) | 0.90 (0.56) | 0.052 |
| Mean number of people living in the same house | 4.9 (2.9) | 13.8 (5.9) | <0.001 |
| **Education level** |  |  | 0.5 |
| Elementary | 1 (10%) | 2 (20%) |  |
| Other | 0 (0%) | 1 (10%) |  |
| Secondary | 2 (20%) | 3 (30%) |  |
| Uninstructed | 1 (10%) | 2 (20%) |  |
| University | 6 (60%) | 2 (20%) |  |
| **Housing type** |  |  | <0.001 |
| Appartement | 9 (90%) | 1 (10%) |  |
| House with concessions | 1 (10%) | 9 (90%) |  |
| **Floor coverings** |  |  | <0.001 |
| No | 0 (0%) | 7 (70%) |  |
| Partial | 0 (0%) | 2 (20%) |  |
| Yes | 10 (100%) | 1 (10%) |  |
| **Wall coverings** |  |  | <0.001 |
| No | 0 (0%) | 7 (70%) |  |
| Partial | 0 (0%) | 1 (10%) |  |
| Yes | 10 (100%) | 2 (20%) |  |
| **Frequency of Incomes** |  |  | 0.034 |
| irregular | 4 (40%) | 6 (100%) |  |
| regular | 6 (60%) | 0 (0%) |  |
| Missing | 0 | 4 |  |

| Characteristic | Residence |  | p-value<sup>2</sup> |
| --- | --- | --- | --- |
|  | High SES<br>N = 10<sup>1</sup> | Low SES<br>N = 10<sup>1</sup> |  |
| **Contact with animals** | 1 (10%) | 6 (100%) | <0.001 |
| Missing | 0 | 4 |  |
| **Attitude when sick** |  |  | 0.057 |
| automedication | 6 (60%) | 1 (10%) |  |
| see a doctor | 4 (40%) | 7 (70%) |  |
| wait to feel better | 0 (0%) | 2 (20%) |  |

<sup>1</sup>n (%); Mean (Min); Mean (SD)
<sup>2</sup>Fisher's exact test; Wilcoxon rank sum test; Pearson's Chi-squared test

**Table S7.** Targeted B-cell panel for validation of SES related surface markers.

| Specificity | Fluorochrome | Company | Catalogue № | Clone | Dilution |
| --- | --- | --- | --- | --- | --- |
| BD Stain Plus |  | BD | 566385 |  | 10 |
| Viability | Live dead blue | Invitrogen | L34962 |  | 500 |
| CD360..IL.21R. | APC | Biolegend | 376105 | W18100A | 20 |
| CD200( OX2) | BV480 | BD | 746397 | MRC OX-104 | 50 |
| CD85j (ILT2) | BUV737 | Biolegend | 310730 | GHI/75 | 50 |
| CD193 (CCR3) | BV711 | BD | 748455 | 5E8 | 50 |
| CD45RB | AF700 | Thermo Fisher | MA5-28589 | MEM-55 | 50 |
| CD166 | BUV805 | BD | 752526 | 3A6 | 50 |
| TLT.2 | PE | Biolegend | 351104 | MIH61 | 50 |
| CD3 | Spark Blue 574 | Biolegend | 300487 | UCHT1 | 200 |
| CD27 | BV785 | Biolegend | 302831 | O323 | 200 |
| CD38 | APC-Fire 810 | Biolegend | 303549 | HIT2 | 200 |
| HLA-DR | BUV496 | BD | 753685 | L243 | 200 |
| CD31 | StarBright V440 | Bio-Rad | MCA1738SBV440 | WM59 | 400 |
| CD185 (CXCR5) | BUV395 | BD | 740266 | RF8B2 | 400 |
| IgG | BV510 | BD | 563247 | G18-145 | 400 |
| CD11c | BV650 | Biolegend | 337237 | Bu15 | 800 |
| CD21 | FITC | Biolegend | 354909 | Bu32 | 800 |
| CD79b..Igβ | BUV615 | BD | 751261 | 3A2-2E7 | 800 |
| CD95..Fas. | PE-Cy5 | Biolegend | 305610 | DX2 | 800 |
| CD55 | BUV563 | BD | 750099 | IA10 | 1600 |
| CD19 | PE-Cy5.5 | eBioscience™ | MHCD1918 | SJ25 | 1600 |
| IgD | RB744 | BD | 758125 | IA6-2 | 1600 |
| IgA | APC-Vio 770 | Miltenyi | 130-113-473 | IS11-8E10 | 1600 |
| CD35 | RB705 | BD | 757700 | E11 | 3200 |
| IgM | Spark Blue 550 | Biolegend | 314555 | MHM-88 | 3200 |
| CD82 | PE-Cy7 | Biolegend | 342109 | ASL-24 | 6400 |

**Table S8.** Targeted myeloid panel for validation of SES related surface markers.

| Specificity | Fluorochrome | Company | Catalogue № | Clone | Dilution |
| --- | --- | --- | --- | --- | --- |
| BD Stain Plus |  | BD | 566385 |  | 10 |
| Viability | Live dead blue | Invitrogen | L34962 |  | 500 |
| CD191 (CCR1) | FITC | Biolegend | 362909 | 5F10B29 | 50 |
| CD182 (CXCR2) | BUV395 | BD | 744201 | 6C6 | 50 |
| CD88..C5aR. | BV786 | BD | 742320 | D53-1473 | 50 |
| CD354 (TREM.1) | BV510 | BD | 743739 | 6B1 | 50 |
| CD155..PVR. | AF700 | Biolegend | 337629 | SKII.4 | 50 |
| CD66 (a,c,e) | BV605 | Biolegend | 342323 | ASL-32 | 50 |
| CD107b (LAMP.2) | Alexa Fluor 647 | BD | 565305 | H4B4 | 100 |
| CD300c | BV480 | BD | 746474 | TX45 | 100 |
| Integrin_β7 | APC-Fire 750 | Biolegend | 321223 | FIB504 | 100 |
| CD100 | BV711 | Biolegend | 745497 | A8 | 200 |
| CD14 | Spark Blue 574 | Biolegend | 325635 | HCD14 | 200 |
| FcεRIα | RB744 | BD | 757890 | AER-37 (CRA-1) | 200 |
| MERTK | PE | Biolegend | 367607 | 590H11G1E3 | 200 |
| CD11c | StarBright V440 | Bio-Rad | MCA2087SBV440 | BU15 | 400 |
| CD163 | PE-Cy5 | Biolegend | 333643 | GHI/61 | 800 |
| CD1c | Super bright 645 | Thermo Fisher | 64-0015-42 | L161 | 800 |
| CD123 | RB780 | BD | 568691 | 7G3 | 800 |
| CD192 (CCR2) | PE-Dazzle 594 | Biolegend | 357221 | K036C2 | 800 |
| HLA-DR | APC-Fire 810 | Biolegend | 307673 | L243 | 800 |
| CD16 | BUV737 | BD | 612786 | 3G8 | 800 |
| CD7 | BUV805 | BD | 742002 | M-T701 | 800 |
| CD180..RP105. | PE-Cy7 | Biolegend | 312909 | MHR73-11 | 3200 |
| CD50 (ICAM.3) | RB705 | BD | 757623 | TU41 | 6400 |

**Table S9.**
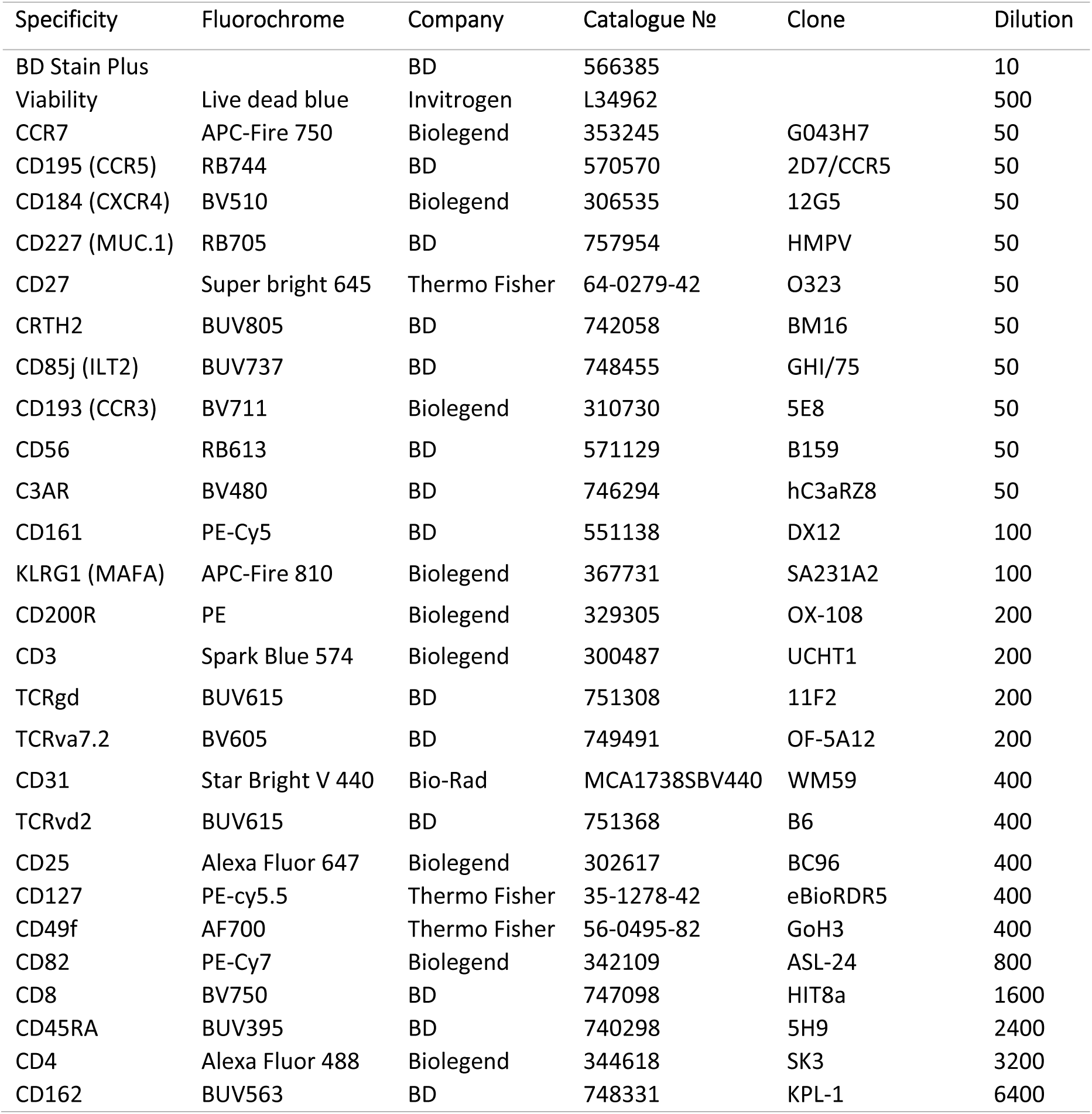
Targeted T-cell panel for validation of SES related surface markers.

| Specificity | Fluorochrome | Company | Catalogue № | Clone | Dilution |
| --- | --- | --- | --- | --- | --- |
| BD Stain Plus |  | BD | 566385 |  | 10 |
| Viability | Live dead blue | Invitrogen | L34962 |  | 500 |
| CCR7 | APC-Fire 750 | Biolegend | 353245 | G043H7 | 50 |
| CD195 (CCR5) | RB744 | BD | 570570 | 2D7/CCR5 | 50 |
| CD184 (CXCR4) | BV510 | Biolegend | 306535 | 12G5 | 50 |
| CD227 (MUC.1) | RB705 | BD | 757954 | HMPV | 50 |
| CD27 | Super bright 645 | Thermo Fisher | 64-0279-42 | O323 | 50 |
| CRTH2 | BUV805 | BD | 742058 | BM16 | 50 |
| CD85j (ILT2) | BUV737 | BD | 748455 | GHI/75 | 50 |
| CD193 (CCR3) | BV711 | Biolegend | 310730 | 5E8 | 50 |
| CD56 | RB613 | BD | 571129 | B159 | 50 |
| C3AR | BV480 | BD | 746294 | hC3aRZ8 | 50 |
| CD161 | PE-Cy5 | BD | 551138 | DX12 | 100 |
| KLRG1 (MAFA) | APC-Fire 810 | Biolegend | 367731 | SA231A2 | 100 |
| CD200R | PE | Biolegend | 329305 | OX-108 | 200 |
| CD3 | Spark Blue 574 | Biolegend | 300487 | UCHT1 | 200 |
| TCRgd | BUV615 | BD | 751308 | 11F2 | 200 |
| TCRva7.2 | BV605 | BD | 749491 | OF-5A12 | 200 |
| CD31 | Star Bright V 440 | Bio-Rad | MCA1738SBV440 | WM59 | 400 |
| TCRvd2 | BUV615 | BD | 751368 | B6 | 400 |
| CD25 | Alexa Fluor 647 | Biolegend | 302617 | BC96 | 400 |
| CD127 | PE-cy5.5 | Thermo Fisher | 35-1278-42 | eBioRDR5 | 400 |
| CD49f | AF700 | Thermo Fisher | 56-0495-82 | GoH3 | 400 |
| CD82 | PE-Cy7 | Biolegend | 342109 | ASL-24 | 800 |
| CD8 | BV750 | BD | 747098 | HIT8a | 1600 |
| CD45RA | BUV395 | BD | 740298 | 5H9 | 2400 |
| CD4 | Alexa Fluor 488 | Biolegend | 344618 | SK3 | 3200 |
| CD162 | BUV563 | BD | 748331 | KPL-1 | 6400 |

**Table S10.** Demography of biosynthesis activity profiling cohort.

| **Characteristic** | **Residence** |  |  |
| --- | --- | --- | --- |
|  | **EU**<br>N = 10<sup>1</sup> | **High SES**<br>N = 10<sup>1</sup> | **Low SES**<br>N = 10<sup>1</sup> |
| **Age, years** | 31 (28 - 40) | 33 (22 - 45) | 31 (19 - 41) |
| **Sex, female** | 5 (50%) | 5 (50%) | 6 (60%) |
| **Height, cm** | NR | 1.75 (0.12) | 1.77 (0.11) |
| *Missing* | 10 | 3 | 3 |
| **Weight, kg** | NR | 74 (15) | 69 (16) |
| *Missing* | 10 | 0 | 0 |
| **BMI, kg/m<sup>2</sup>** | NR | 22.4 (2.5) | 23.9 (6.5) |
| *Missing* | 10 | 3 | 3 |
| **Systolic blood pressure, mmHg** | NR | 130 (120, 130) | 120 (118, 142) |
| *Missing* | 10 | 0 | 0 |
| **Diastolic blood pressure, mmHg** | NR | 80 (80, 90) | 71 (65, 89) |
| *Missing* | 10 | 0 | 0 |
| **Current or past tobacco use** | 0 (0%) | 4 (40%) | 2 (20%) |
| **Passive smoking** | 0 (0%) | 7 (70%) | 3 (30%) |

<sup>1</sup>Median (Min - Max); n (%); Mean (SD); Median (Q1, Q3)

**Table S11.**
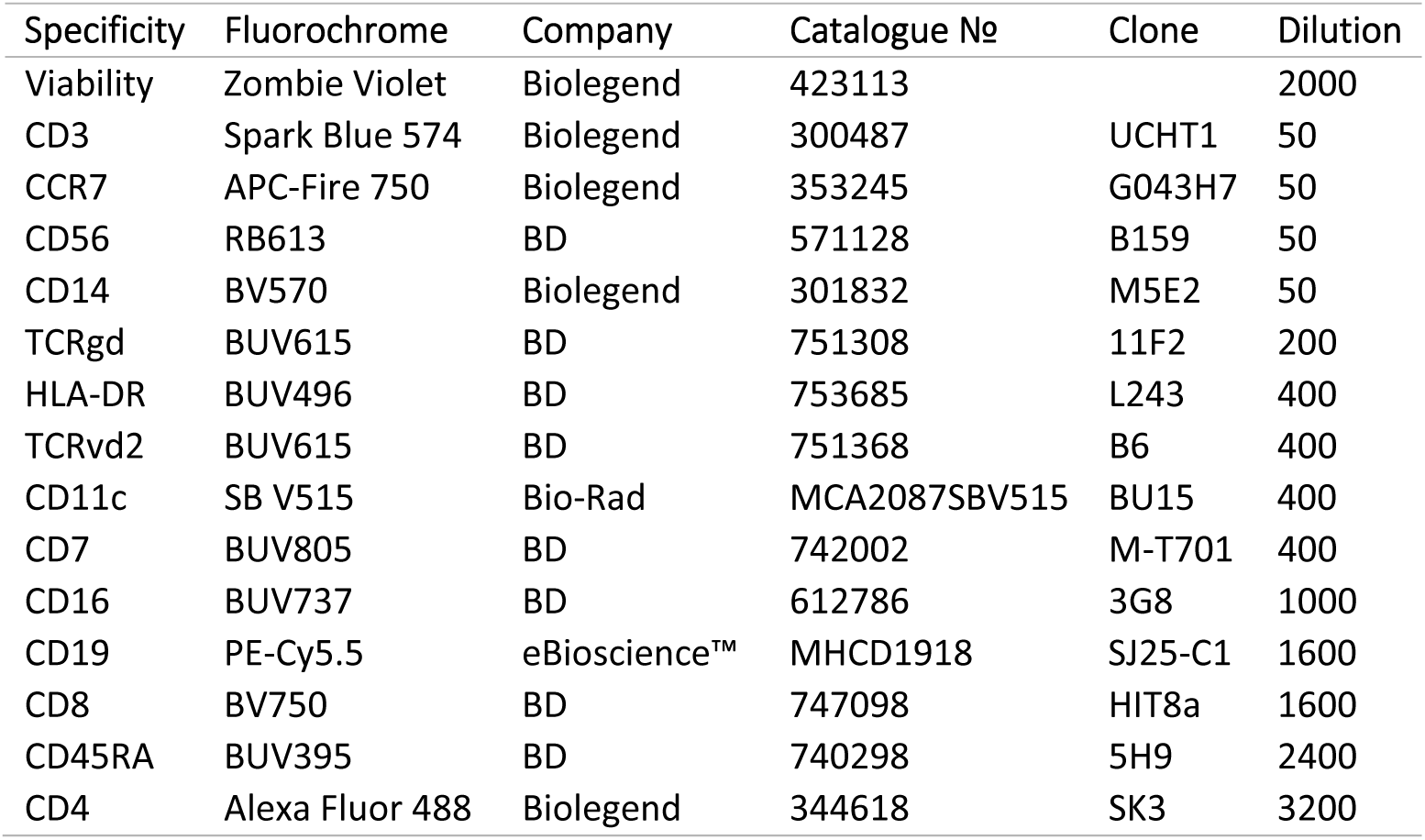
Spectral flow cytometry panel for PBMC biosynthesis activity profiling.

## References

Abavisani, M., Ansari, B., Ebadpour, N., & Sahebkar, A. (2024). How does geographical diversity shape vaccine efficacy? Clin Exp Vaccine Res, 13(4), 271–300. 10.7774/cevr.2024.13.4.271

Amoah, A. S., Agnandji, S. T., Bengtson, M., Downs, J. A., Esen, M., Giddaluru, J., Jochems, S. P., Kaisar, M. M. M., Makinde, J., Manurung, M. D., Mbow, M., Moncunill, G., Murugan, R., Mutai, J., Nakaya, H. I., Nkurunungi, G., Palmblad, M., Pyuza, J. J., Stam, K. A.,…Participants of the Lorentz Center Workshop titled Connecting people to reverse vaccine, h. (2025). Tackling vaccine hyporesponsiveness through global collaboration, diverse population studies, and data integration. Lancet Microbe, 101263. 10.1016/j.lanmic.2025.101263

Azad, M. B., Lissitsyn, Y., Miller, G. E., Becker, A. B., HayGlass, K. T., & Kozyrskyj, A. L. (2012). Influence of socioeconomic status trajectories on innate immune responsiveness in children. PLoS One, 7(6), e38669. 10.1371/journal.pone.0038669

Bertrand, A., Sugrue, J., Lou, T., Bourke, N. M., Quintana-Murci, L., Saint-Andre, V., O’Farrelly, C., Duffy, D., & Milieu Interieur, C. (2024). Impact of socioeconomic status on healthy immune responses in humans. Immunol Cell Biol, 102(7), 618–629. 10.1111/imcb.12789

Brenchley, J. M., Karandikar, N. J., Betts, M. R., Ambrozak, D. R., Hill, B. J., Crotty, L. E., Casazza, J. P., Kuruppu, J., Migueles, S. A., Connors, M., Roederer, M., Douek, D. C., & Koup, R. A. (2003). Expression of CD57 defines replicative senescence and antigen-induced apoptotic death of CD8+ T cells. Blood, 101(7), 2711–2720. 10.1182/blood-2002-07-2103

Brodin, P., & Davis, M. M. (2017). Human immune system variation. Nat Rev Immunol, 17(1), 21–29. 10.1038/nri.2016.125

Brodin, P., Jojic, V., Gao, T., Bhattacharya, S., Angel, C. J., Furman, D., Shen-Orr, S., Dekker, C. L., Swan, G. E., Butte, A. J., Maecker, H. T., & Davis, M. M. (2015). Variation in the human immune system is largely driven by non-heritable influences. Cell, 160(1-2), 37–47. 10.1016/j.cell.2014.12.020

Buck, M. D., O’Sullivan, D., Klein Geltink, R. I., Curtis, J. D., Chang, C. H., Sanin, D. E., Qiu, J., Kretz, O., Braas, D., van der Windt, G. J., Chen, Q., Huang, S. C., O’Neill, C. M., Edelson, B. T., Pearce, E. J., Sesaki, H., Huber, T. B., Rambold, A. S., & Pearce, E. L. (2016). Mitochondrial Dynamics Controls T Cell Fate through Metabolic Programming. Cell, 166(1), 63–76. 10.1016/j.cell.2016.05.035

de Jong, S. E., van Unen, V., Manurung, M. D., Stam, K. A., Goeman, J. J., Jochems, S. P., Hollt, T., Pezzotti, N., Mouwenda, Y. D., Betouke Ongwe, M. E., Lorenz, F. R., Kruize, Y. C. M., Azimi, S., Konig, M. H., Vilanova, A., Eisemann, E., Lelieveldt, B. P. F., Roestenberg, M., Sim, B. K. L.,…Yazdanbakhsh, M. (2021). Systems analysis and controlled malaria infection in Europeans and Africans elucidate naturally acquired immunity. Nat Immunol, 22(5), 654–665. 10.1038/s41590-021-00911-7

Derre, L., Rivals, J. P., Jandus, C., Pastor, S., Rimoldi, D., Romero, P., Michielin, O., Olive, D., & Speiser, D. E. (2010). BTLA mediates inhibition of human tumor-specific CD8+ T cells that can be partially reversed by vaccination. J Clin Invest, 120(1), 157–167. 10.1172/JCI40070

Dotmatics. (2025). OMIQ Data Analysis Software. In Dotmatics. https://www.omiq.ai/

Dowd, J. B., Aiello, A. E., & Alley, D. E. (2009). Socioeconomic disparities in the seroprevalence of cytomegalovirus infection in the US population: NHANES III. Epidemiol Infect, 137(1), 58–65. 10.1017/S0950268808000551

Dunkelberger, J. R., & Song, W. C. (2010). Complement and its role in innate and adaptive immune responses. Cell Res, 20(1), 34–50. 10.1038/cr.2009.139

Evans, G. W., & Kantrowitz, E. (2002). Socioeconomic status and health: the potential role of environmental risk exposure. Annu Rev Public Health, 23, 303–331. 10.1146/annurev.publhealth.23.112001.112349

Ignacio, B. J., Dijkstra, J., Mora, N., Slot, E. F. J., van Weijsten, M. J., Storkebaum, E., Vermeulen, M., & Bonger, K. M. (2023). THRONCAT: metabolic labeling of newly synthesized proteins using a bioorthogonal threonine analog. Nature Communications, 14(1). ARTN 3367 10.1038/s41467-023-39063-7

Jao, C. Y., & Salic, A. (2008). Exploring RNA transcription and turnover in vivo by using click chemistry. Proc Natl Acad Sci U S A, 105(41), 15779–15784. 10.1073/pnas.0808480105

Kim, W., Zhou, J. Q., Horvath, S. C., Schmitz, A. J., Sturtz, A. J., Lei, T., Liu, Z., Kalaidina, E., Thapa, M., Alsoussi, W. B., Haile, A., Klebert, M. K., Suessen, T., Parra-Rodriguez, L., Mudd, P. A., Whelan, S. P. J., Middleton, W. D., Teefey, S. A., Pusic, I.,…Ellebedy, A. H. (2022). Germinal centre-driven maturation of B cell response to mRNA vaccination. Nature, 604(7904), 141–145. 10.1038/s41586-022-04527-1

Klopack, E. T., Crimmins, E. M., Cole, S. W., Seeman, T. E., & Carroll, J. E. (2022). Social stressors associated with age-related T lymphocyte percentages in older US adults: Evidence from the US Health and Retirement Study. Proc Natl Acad Sci U S A, 119(25), e2202780119. 10.1073/pnas.2202780119

Kuchen, S., Robbins, R., Sims, G. P., Sheng, C., Phillips, T. M., Lipsky, P. E., & Ettinger, R. (2007). Essential role of IL-21 in B cell activation, expansion, and plasma cell generation during CD4+ T cell-B cell collaboration. J Immunol, 179(9), 5886–5896. 10.4049/jimmunol.179.9.5886

Kurtovic, L., & Beeson, J. G. (2021). Complement Factors in COVID-19 Therapeutics and Vaccines. Trends Immunol, 42(2), 94–103. 10.1016/j.it.2020.12.002

Linterman, M. A., Beaton, L., Yu, D., Ramiscal, R. R., Srivastava, M., Hogan, J. J., Verma, N. K., Smyth, M. J., Rigby, R. J., & Vinuesa, C. G. (2010). IL-21 acts directly on B cells to regulate Bcl-6 expression and germinal center responses. J Exp Med, 207(2), 353–363. 10.1084/jem.20091738

Liu, Z., Wang, H., Li, Z., Dress, R. J., Zhu, Y., Zhang, S., De Feo, D., Kong, W. T., Cai, P., Shin, A., Piot, C., Yu, J., Gu, Y., Zhang, M., Gao, C., Chen, L., Wang, H., Vetillard, M., Guermonprez, P.,…Ginhoux, F. (2023). Dendritic cell type 3 arises from Ly6C(+) monocyte-dendritic cell progenitors. Immunity, 56(8), 1761–1777 e1766. 10.1016/j.immuni.2023.07.001

MacGillivray, D. M., & Kollmann, T. R. (2014). The role of environmental factors in modulating immune responses in early life. Front Immunol, 5, 434. 10.3389/fimmu.2014.00434

Magill, L., Adriani, M., Berthou, V., Chen, K., Gleizes, A., Hacein-Bey-Abina, S., Hincelin-Mery, A., Mariette, X., Pallardy, M., Spindeldreher, S., Szely, N., Isenberg, D. A., Manson, J. J., Jury, E. C., & Mauri, C. (2018). Low Percentage of Signal Regulatory Protein alpha/beta(+) Memory B Cells in Blood Predicts Development of Anti-drug Antibodies (ADA) in Adalimumab-Treated Rheumatoid Arthritis Patients. Front Immunol, 9, 2865. 10.3389/fimmu.2018.02865

Manurung, M. D., Heieis, G. A., Konig, M., Azimi, S., Ndao, M., Veldhuizen, T., Hoving, D., Hoekstra, P. T., Kruize, Y. C. M., Wammes, L. J., Menafra, R., Cisse, M., Mboup, S., Dieye, A., Kloet, S., Tahapary, D. L., Supali, T., Wuhrer, M., Hokke, C. H.,…Mbow, M. (2025). Systems analysis unravels a common rural-urban gradient in immunological profile, function, and metabolic dependencies. Sci Adv, 11(18), eadu0419. 10.1126/sciadv.adu0419

Mbow, M., de Jong, S. E., Meurs, L., Mboup, S., Dieye, T. N., Polman, K., & Yazdanbakhsh, M. (2014). Changes in immunological profile as a function of urbanization and lifestyle. Immunology, 143(4), 569–577. 10.1111/imm.12335

Mbow, M., Hoving, D., Cisse, M., Diallo, I., Honkpehedji, Y. J., Huisman, W., Pothast, C. R., Jongsma, M. L. M., Konig, M. H., de Kroon, A. C., Linh, L. T. K., Azimi, S., Tak, T., Kruize, Y. C. M., Kurniawan, F., Dia, Y. A., Zhang, J. L. H., Prins, C., Roukens, A. H. E.,…team, C.-L. U. M. C. s. (2025). Immune responses to SARS-CoV-2 in sub-Saharan Africa and western Europe: a retrospective, population-based, cross-sectional study. Lancet Microbe, 6(2), 100942. 10.1016/j.lanmic.2024.07.005

McInnes, L., & Healy, J. (2018). UMAP: Uniform Manifold Approximation and Projection for Dimension Reduction. 10.48550/arXiv.1802.03426

O’Neill, L. A., Kishton, R. J., & Rathmell, J. (2016). A guide to immunometabolism for immunologists. Nat Rev Immunol, 16(9), 553–565. 10.1038/nri.2016.70

Pawelec, G. (2017). Immunosenescence and cancer. Biogerontology, 18(4), 717–721. 10.1007/s10522-017-9682-z

Posit team. (2025). RStudio: Integrated Development Environment for R. In Posit Software, PBC. http://www.posit.co/

Potaczek, D. P., & Kabesch, M. (2012). Current concepts of IgE regulation and impact of genetic determinants. Clin Exp Allergy, 42(6), 852–871. 10.1111/j.1365-2222.2011.03953.x

Pulendran, B., & Ahmed, R. (2011). Immunological mechanisms of vaccination. Nat Immunol, 12(6), 509–517. 10.1038/ni.2039

Pyuza, J. J., van Dorst, M., Stam, K., Wammes, L., Konig, M., Kullaya, V. I., Kruize, Y., Huisman, W., Andongolile, N., Ngowi, A., Shao, E. R., Mremi, A., Hogendoorn, P. C. W., Msuya, S. E., Jochems, S. P., de Steenhuijsen Piters, W. A. A., & Yazdanbakhsh, M. (2024). Lifestyle score is associated with cellular immune profiles in healthy Tanzanian adults. Brain Behav Immun Health, 41, 100863. 10.1016/j.bbih.2024.100863

R Core Team. (2025). R: A Language and Environment for Statistical Computing. In R Foundation for Statistical Computing. https://www.R-project.org/

Ravi, S., Shanahan, M. J., Levitt, B., Harris, K. M., & Cole, S. W. (2024). Socioeconomic inequalities in early adulthood disrupt the immune transcriptomic landscape via upstream regulators. Sci Rep, 14(1), 1255. 10.1038/s41598-024-51517-6

Relman, D. A., & Lipsitch, M. (2018). Microbiome as a tool and a target in the effort to address antimicrobial resistance. Proc Natl Acad Sci U S A, 115(51), 12902–12910. 10.1073/pnas.1717163115

Ronsmans, S., Sorig Hougaard, K., Nawrot, T. S., Plusquin, M., Huaux, F., Jesus Cruz, M., Moldovan, H., Verpaele, S., Jayapala, M., Tunney, M., Humblet-Baron, S., Dirven, H., Cecilie Nygaard, U., Lindeman, B., Duale, N., Liston, A., Meulengracht Flachs, E., Kastaniegaard, K., Ketzel, M.,…Hoet, P. H. M. (2022). The EXIMIOUS project-Mapping exposure-induced immune effects: connecting the exposome and the immunome. Environ Epidemiol, 6(1), e193. 10.1097/EE9.0000000000000193

Rook, G. A. W. (2022). Evolution, the Immune System, and the Health Consequences of Socioeconomic Inequality. mSystems, 7(2), e0143821. 10.1128/msystems.01438-21

Santiago, H. C., Bennuru, S., Boyd, A., Eberhard, M., & Nutman, T. B. (2011). Structural and immunologic cross-reactivity among filarial and mite tropomyosin: implications for the hygiene hypothesis. J Allergy Clin Immunol, 127(2), 479–486. 10.1016/j.jaci.2010.11.007

Snyder-Mackler, N., Sanz, J., Kohn, J. N., Brinkworth, J. F., Morrow, S., Shaver, A. O., Grenier, J. C., Pique-Regi, R., Johnson, Z. P., Wilson, M. E., Barreiro, L. B., & Tung, J. (2016). Social status alters immune regulation and response to infection in macaques. Science, 354(6315), 1041–1045. 10.1126/science.aah3580

Stringhini, S., Polidoro, S., Sacerdote, C., Kelly, R. S., van Veldhoven, K., Agnoli, C., Grioni, S., Tumino, R., Giurdanella, M. C., Panico, S., Mattiello, A., Palli, D., Masala, G., Gallo, V., Castagne, R., Paccaud, F., Campanella, G., Chadeau-Hyam, M., & Vineis, P. (2015). Life-course socioeconomic status and DNA methylation of genes regulating inflammation. Int J Epidemiol, 44(4), 1320–1330. 10.1093/ije/dyv060

Tehranifar, P., Wu, H. C., Fan, X., Flom, J. D., Ferris, J. S., Cho, Y. H., Gonzalez, K., Santella, R. M., & Terry, M. B. (2013). Early life socioeconomic factors and genomic DNA methylation in mid-life. Epigenetics, 8(1), 23–27. 10.4161/epi.22989

Turner, J. S., Zhou, J. Q., Han, J., Schmitz, A. J., Rizk, A. A., Alsoussi, W. B., Lei, T., Amor, M., McIntire, K. M., Meade, P., Strohmeier, S., Brent, R. I., Richey, S. T., Haile, A., Yang, Y. R., Klebert, M. K., Suessen, T., Teefey, S., Presti, R. M.,…Ellebedy, A. H. (2020). Human germinal centres engage memory and naive B cells after influenza vaccination. Nature, 586(7827), 127–132. 10.1038/s41586-020-2711-0

Van’t Sant, L. J., White, J. J., Hoeijmakers, J. H. J., Vermeij, W. P., & Jaarsma, D. (2021). In vivo 5-ethynyluridine (EU) labelling detects reduced transcription in Purkinje cell degeneration mouse mutants, but can itself induce neurodegeneration. Acta Neuropathol Commun, 9(1), 94. 10.1186/s40478-021-01200-y

van den Beukel, M. D., Zhang, L., van der Meulen, S., Borggreven, N. V., Nugteren, S., Brouwer, M. C., Pouw, R. B., Gelderman, K. A., de Ru, A. H., Janssen, G. M. C., van Veelen, P. A., Knevel, R., Parren, P., & Trouw, L. A. (2025). Post-translationally modified proteins bind and activate complement with implications for cellular uptake and autoantibody formation. J Autoimmun, 155, 103444. 10.1016/j.jaut.2025.103444

van Dorst, M., Azimi, S., Wahyuni, S., Amaruddin, A. I., Sartono, E., Wammes, L. J., Yazdanbakhsh, M., & Jochems, S. P. (2022). Differences in Bacterial Colonization and Mucosal Responses Between High and Low SES Children in Indonesia. Pediatr Infect Dis J, 41(6), 496–506. 10.1097/INF.0000000000003525

van Dorst, M., Pyuza, J. J., Nkurunungi, G., Kullaya, V. I., Smits, H. H., Hogendoorn, P. C. W., Wammes, L. J., Everts, B., Elliott, A. M., Jochems, S. P., & Yazdanbakhsh, M. (2024). Immunological factors linked to geographical variation in vaccine responses. Nat Rev Immunol, 24(4), 250–263. 10.1038/s41577-023-00941-2

Van Gassen, S., Callebaut, B., Van Helden, M. J., Lambrecht, B. N., Demeester, P., Dhaene, T., & Saeys, Y. (2015). FlowSOM: Using self-organizing maps for visualization and interpretation of cytometry data. Cytometry A, 87(7), 636–645. 10.1002/cyto.a.22625

Van Gassen, S., Gaudilliere, B., Angst, M. S., Saeys, Y., & Aghaeepour, N. (2020). CytoNorm: A Normalization Algorithm for Cytometry Data. Cytometry Part A, 97(3), 268–278. 10.1002/cyto.a.23904

Vrieling, F., van der Zande, H. J. P., Naus, B., Smeehuijzen, L., van Heck, J. I. P., Ignacio, B. J., Bonger, K. M., van den Bossche, J., Kersten, S., & Stienstra, R. (2024). CENCAT enables immunometabolic profiling by measuring protein synthesis via bioorthogonal noncanonical amino acid tagging. Cell Reports Methods, 4(10). ARTN 100883 10.1016/j.crmeth.2024.100883

Zheng, T., Ding, C., Lai, S., Gao, Y., Lyu, C., Liu, C., Shi, J., Li, X., Li, M., Meng, H., Li, M., Liang, Y., Tai, S., Cheng, L., Zhang, Y., Li, L., Han, P., Sun, B., Liu, T.,…Zhang, X. (2025). CD160 dictates anti-PD-1 immunotherapy resistance by regulating CD8(+) T cell exhaustion in colorectal cancer. Nat Cell Biol, 27(9), 1555–1571. 10.1038/s41556-025-01753-3

Zotos, D., Coquet, J. M., Zhang, Y., Light, A., D’Costa, K., Kallies, A., Corcoran, L. M., Godfrey, D. I., Toellner, K. M., Smyth, M. J., Nutt, S. L., & Tarlinton, D. M. (2010). IL-21 regulates germinal center B cell differentiation and proliferation through a B cell-intrinsic mechanism. J Exp Med, 207(2), 365–378. 10.1084/jem.20091777

